# Secretion and signaling properties of Wnt11 and Wnt3 are determined by their N-terminus

**DOI:** 10.1101/2025.07.02.662750

**Authors:** Oksana Voloshanenko, Nachiket Dhonnar, Diego Aponte-Santamaría, Carina Seidl, Lucie Wolf, Dominique Kranz, Alena Ivanova, Eddye J. Figueroa Marquez, Bianca Pokrandt, Iris Augustin, Irmgard M. Sinning, Port Fillip, Christof Niehrs, Britta Brügger, Michael Boutros

## Abstract

Wnt proteins are evolutionarily conserved signaling molecules that control cell-cell communication in health and disease. While different Wnt proteins share a requirement for palmitoleoylation and the cargo receptor Evi/Wls for their secretion, they interact with diverse signaling receptors and co-receptors to elicit distinct outcomes in signal-receiving cells. The molecular mechanisms underlying this contrast between conserved post-translational modification and secretion and diverse receptor interactions remain poorly understood. Here, we demonstrate that Wnt11 maintains partial secretion capability in the absence of Evi/Wls. Unexpectedly, the secreted protein is also non-palmitoleoylated, yet retains its ability to activate downstream signaling in receiving cells through the Frizzled 6 receptor. We identify the N-terminus as the mediator of this unusual behavior, as Wnt3, which strictly requires Evi/Wls, becomes Evi/Wls-independent when its N-terminus is replaced with that of Wnt11. In different cell lines and in *Xenopus laevis* embryos, the chimeric Wnt11^Nterm^-Wnt3 protein recapitulates known Wnt11 phenotypes, demonstrating that the N-terminal portion of Wnt11 is sufficient to determine signaling specificity. Our findings reveal novel determinants of Wnt protein specificity and provide mechanistic insights into the molecular basis of Wnt secretion and downstream signaling.

## INTRODUCTION

Wnt proteins are essential regulators of development and adult homeostasis, and dysregulation of Wnt signaling has been implicated in numerous human diseases (Niehrs, 2012; Nusse & Clevers, 2017; Zhan *et al*, 2017). Wnt ligands are a family of secreted lipid-modified glycoproteins that are evolutionarily conserved throughout the animal kingdom, from the sea anemone *Nematostella* to humans (Holzem *et al*, 2024; Holstein, 2012). Most mammalian genomes encode 19 Wnt proteins, which elicit context-dependent responses through their interactions with diverse cell surface receptor complexes (Boutros & Mlodzik, 1999; Boutros *et al*, 1998; Niehrs, 2012).

The secretion of Wnt proteins from signal-sending cells is tightly regulated (Takada *et al*, 2017; Gross & Boutros, 2013; Herr *et al*, 2012; Wolf & Boutros, 2023; Maurice & Angers, 2025). Following translation into the endoplasmic reticulum, Wnts are lipid-modified by the membrane-bound O-acyltransferase porcupine (PORCN) at a conserved serine residue (Ser209 in Wnt3a) (Takada *et al*, 2006). Multiple lines of evidence suggest that palmitoleoylation is essential for Wnt secretion, as mutations (Wang *et al*, 2007; Grzeschik *et al*, 2007), silencing (Kadowaki *et al*, 1996; Takada *et al*, 2006), chemical inhibition, or knock-out (Barrott *et al*, 2011; Biechele *et al*, 2011) of PORCN inhibit secretion of multiple Wnt ligands, including Wnt1, Wnt2, Wnt3/3a, Wnt5a, Wnt6, Wnt7a and Wnt9a (Liu *et al*, 2022, 2013; Dodge *et al*, 2012). Palmitoleoylated Wnts interact with the multispan transmembrane cargo receptor Evi/Wls/GPR177, which is conserved across metazoa and is required for Wnt transport and secretion (Goodman *et al*, 2006; Bartscherer *et al*, 2006; Bänziger *et al*, 2006). PORCN-mediated acylation appears necessary for this interaction (Coombs *et al*, 2010; Herr & Basler, 2012), consistent with recent cryo-EM structures revealing a central cavity in Evi/Wls that accommodates the Wnt protein’s palmitoleate chain (Zhong *et al*, 2021; Nygaard *et al*, 2021). *Evi/Wls*-deficient embryos phenocopy *Wnt1* and *Wnt3* knockout in mice, and tissue-specific disruption of Evi/Wls impairs Wnt signaling, supporting the fundamental importance of Evi/Wls for Wnt secretion (Korn *et al*, 2014; Augustin *et al*, 2013; Carpenter *et al*, 2010; Fu *et al*, 2011). While these findings suggest that mammalian Wnt proteins generally depend on a conserved secretory machinery centered around PORCN and Evi/Wls, recent studies reveal potential exceptions. For example, *Xenopus* Wnt8 and human Wnt7a retain functionality without acylation and can be secreted independently of PORCN, indicating that Wnt secretion mechanisms might be more diverse than so far anticipated (Gurriaran-Rodriguez *et al*, 2024; Speer *et al*, 2019; von Maltzahn *et al*, 2013).

Secreted Wnt proteins act on receiving cells to regulate a wide range of cellular functions, including stemness, differentiation, proliferation, migration and genome integrity (Nusse & Clevers, 2017; Hayat *et al*, 2022). Canonical Wnt signaling activates the transcriptional co-activator β-catenin to regulate target gene expression, while non-canonical signaling involves diverse post-transcriptional mechanisms to alter cellular functions. Mechanistically, Wnts initiate downstream signaling through binding to frizzled (FZD) receptors and various co-receptors. Multiple factors are believed to determine downstream signaling outcomes, including the specific FZD receptors engaged by each Wnt, the strength of receptor activation, and the cellular state of the receiving cell (Dijksterhuis *et al*, 2015). Among the 10 FZD receptors, FZD3 and FZD6 are structurally and functionally distinct from other FZDs. These receptors exhibit traditional GPCR activity and primarily induce non-canonical/planar cell polarity (PCP) signaling in the nervous system and during development (Kozielewicz *et al*, 2020; Zheng & Sheng, 2024).

A key outstanding question is what determines the signaling specificity of different Wnt ligands. One possibility is that the specificity is intrinsic to each Wnt protein. For example, Wnt1, Wnt3 and Wnt3a have so far been described to exclusively induce canonical signaling in different contexts, while Wnt5a, Wnt5b and Wnt11 signal through non-canonical, β-catenin independent pathways. Studies using chimeric Wnt proteins in *Xenopus* have revealed that signaling specificity can be transplanted between Wnts by domain swaps. For example, canonical signaling activity appears to be encoded in the C-terminal region of Wnt8 (Du *et al*, 1995; Wallkamm *et al*, 2016). Wnt7 ligands have recently been shown to recruit specific co-receptors through N-terminal residues not found in any of the other Wnt ligands (Qi *et al*, 2023) and the canonical Wnt3a and the non-canonical Wnt11 ligands are differentially glycosylated at their N-termini and secreted preferentially from the basolateral or apical side of polarized cells, respectively (Yamamoto *et al*, 2013). However, in other instances, the same Wnt protein can induce canonical or non-canonical signaling in receiving cells. For example, Wnt4, Wnt9a, and Wnt9b have all been shown to be able to induce β-catenin-dependent or -independent signaling in a context-dependent manner (Ide & Grainger, 2024; Zhang *et al*, 2021).

Here, we examine key molecular features that determine Wnt signaling specificity and their influence on both protein secretion and signal transduction. We found that Wnt11 can, in part, be secreted independently of Evi/Wls, while Wnt5a is strictly dependent on Evi/Wls. The Wnt11 protein that is secreted in the absence of Evi/Wls is not palmitoleoylated, yet retains signaling activity. We identified the N-terminus, and more specifically Asn40, as the key regulator of this unique behavior. A chimeric Wnt11^1-48^-Wnt3 is secreted independently of Evi/Wls, demonstrating that this peptide is sufficient to render Wnts independent of their cargo receptor. Intriguingly, also Wnt3 downstream signaling specificity was changed from canonical to non-canonical signaling through exchange of the N-terminal 48 amino acids. Wnt11^1-48^-Wnt3 mediates non-canonical Wnt signaling, increases invasion of melanoma cells and induces defects in convergent extension movement in *Xenopus*. Together, our data indicate that the N-terminus of at least some Wnt proteins is necessary and sufficient to determine signaling specificity and dependence on the conserved Wnt secretion machinery.

## RESULTS

### Wnt11 can be secreted independent of the Wnt cargo receptor Evi/Wls

To study secretion and signaling of Wnt proteins that elicit non-canonical signaling in signal receiving cells, we developed an *in vitro* assay using cultured mammalian cells. The assay is based on previous observations that non-canonical Wnt proteins can inhibit β-catenin-dependent signaling induced by canonical Wnts (Westfall *et al*, 2003; Topol *et al*, 2003). We transfected human embryonic kidney 293T (HEK293T) cells with a widely used reporter system for β-catenin-dependent signaling consisting of TCF4/Wnt and actin-*Renilla* plasmids, and co-expressed either Wnt5a or Wnt11. We then activated β-catenin-dependent signaling by treating cells with recombinant Wnt3a (Figure 1A). Addition of recombinant Wnt3a protein alone (Ctrl condition) resulted in a concentration-dependent increase in TCF4/Wnt reporter activity. Expression of either Wnt5a or Wnt11 with addition of recombinant Wnt3a did not activate the TCF4/Wnt reporter (Figure 1B). Importantly, co-expression of Wnt5a or Wnt11 strongly inhibited TCF4/Wnt-reporter activation in response to recombinant Wnt3a, indicating that this assay effectively reports on signaling activity of non-canonical Wnt proteins (Figure 1B).

**Figure 1.**
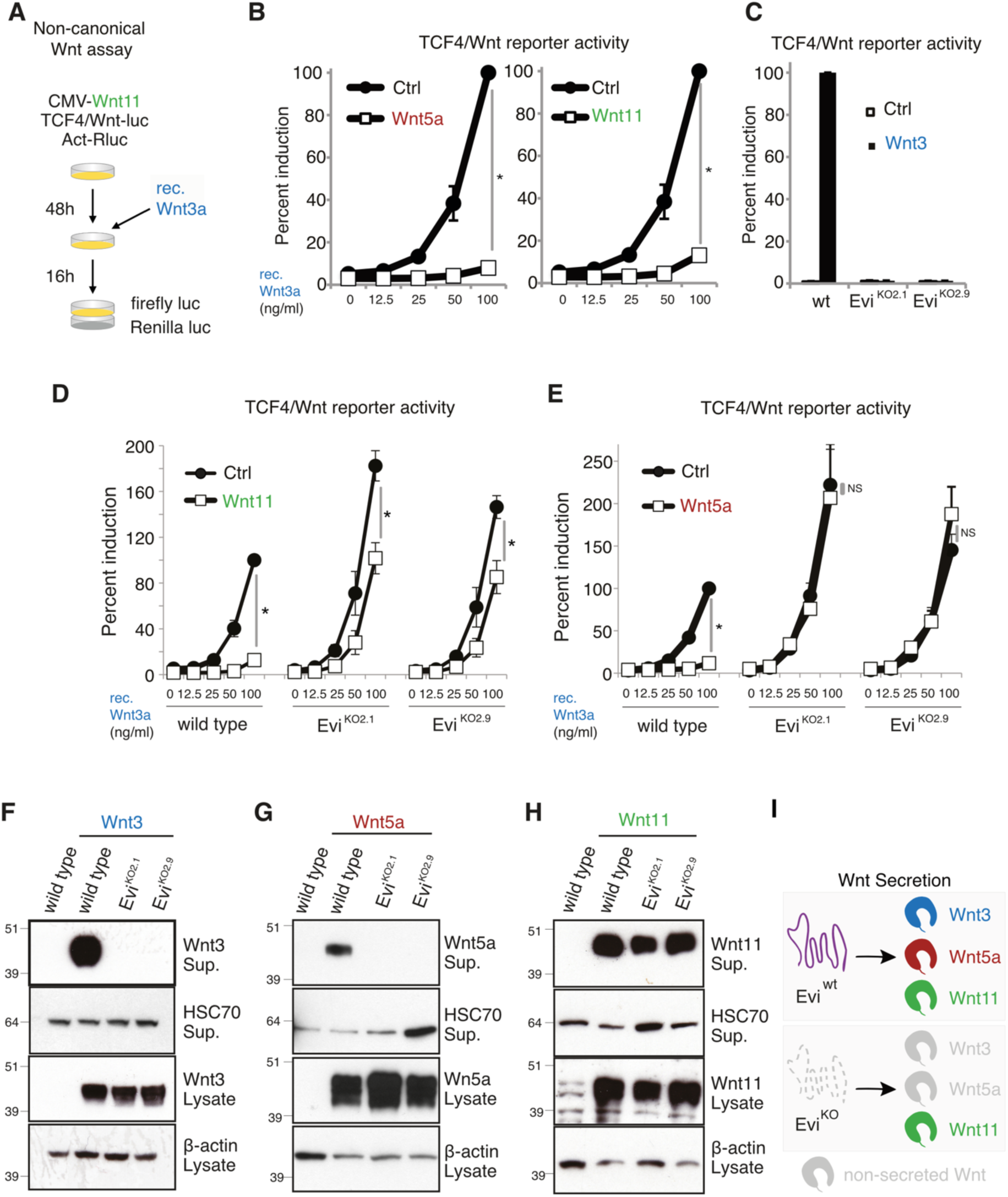
Functional Wnt11 is secreted from Evi knockout cells. **A)** Schematic representation of the non-canonical Wnt assay. **B)** Wnt5a and Wnt11 block β-catenin-dependent Wnt signaling. **C)** Wnt3 expressed in Evi^KO^ cells does not induce β-catenin-dependent signaling. **D,E)** Wnt11 expression in Evi^KO^ cells reduces recombinant Wnt3a-induced TCF4/Wnt-reporter activity, while Wnt5a expressed in Evi^KO^ cells does not inhibit reporter activity. B-E) Indicated HEK293T cells were transfected with Wnt11, Wnt5a or Wnt3, TCF4/Wnt-firefly luciferase and actin-*Renilla* reporters for 48 hrs. B,D,E) If indicated recombinant mouse Wnt3a, in the specified amounts, was added 16 hours before the readout to activate β-catenin-dependent signaling. B-E) Results of 4 independent experiments are shown as mean ± SEM. * *p<0,05, NS -* non-significant. Unpaired *t* test with Welch’s correction. Exact *p* values are shown in Table S2. **F,G)** Wnt3 and Wnt5a secretion is blocked in HEK293T Evi^KO^ cells. **H)** Wnt11 is secreted from Evi^KO^ HEK293T cells. F-H) Indicated cells were transfected with the Wnt3, Wnt5a or Wnt11 plasmids. 48 hrs later supernatants and lysates of cells were collected. Wnts were precipitated from the supernatant (Sup.) with the help of Blue sepharose. β-actin and HSC70 served as the loading control for lysate and supernatant, respectively. Western blots are shown as representatives of 3 independent experiments. **I)** Schematic representation of Wnt secretion and their dependence on Evi/Wls.

Next, we used this assay to test for dependence of non-canonical Wnt proteins on the cargo-receptor Evi/Wls. To this end, we performed it in parallel in HEK293T wild type and Evi/Wls knock-out (Evi^KO^) cell lines previously generated by CRISPR-Cas9 genome editing (Glaeser *et al*, 2018). We used independent single cell clones with frameshift mutations in Evi/Wls (Table S1), which lack Evi/Wls protein (Figure S1A,B) and are unable to secrete functional Wnt3. As a result, they fail to induce the TCF4/Wnt reporter through co-expressed Wnt3 (Figure 1C). Surprisingly, when using recombinant Wnt3a to induce Wnt signaling, Wnt11 mediated substantial inhibition of TCF4/Wnt reporter also in the absence of Evi/Wls (Figure 1D). In contrast, Wnt5a was no longer able to interfere with recombinant Wnt3a signaling in Evi^KO^ cells (Figure 1E). These results could indicate that Wnt11-mediated inhibition of canonical Wnt signaling occurs cell autonomously or that Wnt11 is still secreted in the absence of Evi/Wls. We therefore tested whether Wnt11 can be detected in the supernatant of Evi^KO^ cells. We found that secretion of Wnt3 and Wnt5a was completely blocked in the absence of Evi/Wls (Figure 1F,G,I), whereas Wnt11 protein was still detectable in the supernatant (Figure 1H,I).

To confirm that these findings do not represent artifacts of single cell clone generation, sgRNA off-target effects, or mutations specific to exon 3 of *Evi/Wls*, we validated our results using an independent clone generated with a sgRNA targeting exon 2 (Table S1, Figure S1A,C). To further exclude potential compensation by unknown Evi/Wls splice isoforms, we also generated HEK293T cell pools with mutations in exon 1 (Figure S1A). In these pools, Wnt5a was undetectable in the supernatant, while Wnt11 was still secreted, albeit at reduced levels (Figure S1D). Additionally, we examined Wnt11 secretion from both Evi^KO^ and wild type cells transfected with decreasing amounts of Wnt11 plasmid. These experiments revealed that Evi^KO^ cells consistently secrete less Wnt11 compared to wild type controls, with this reduction remaining proportionally constant across all expression levels (Figure S1C). Collectively, these results confirm that the observed Wnt11 secretion in the absence of Evi/Wls is not due to clonal artifacts, sgRNA off-target effects, or compensation through alternative splicing.

We next investigated whether Wnt11 secretion occurred through an active cellular export mechanism. Treatment with Brefeldin A, a general inhibitor of conventional secretion *via* the secretory pathway, blocked Evi/Wls-independent Wnt11 secretion. This confirms that Wnt11 was actively exported through a vesicular secretion route rather than through cell death or membrane disruption (Figure S1E). Supporting this conclusion, FACS analysis revealed that Wnt11 was present on the cell surface of Evi^KO^ cells, while Wnt5a was not detectable (Figure S2A).

Next, we tested whether Wnt11 can also be secreted independently of the Wnt cargo-receptor in other cellular contexts. We generated HCT116 Evi^KO^ cells, which endogenously express Wnt11 and Wnt10b (Figure 2A, Table S1) (Voloshanenko *et al*, 2013). Wnt11 remained secreted in the absence of Evi/Wls, although at reduced levels compared to wild type cells, whereas Wnt10b secretion was completely blocked (Figure 2A). Knockdown with four independent siRNAs confirmed the specificity of the Wnt11 antibody towards endogenous Wnt11 (Figure S2B).

**Figure 2.**
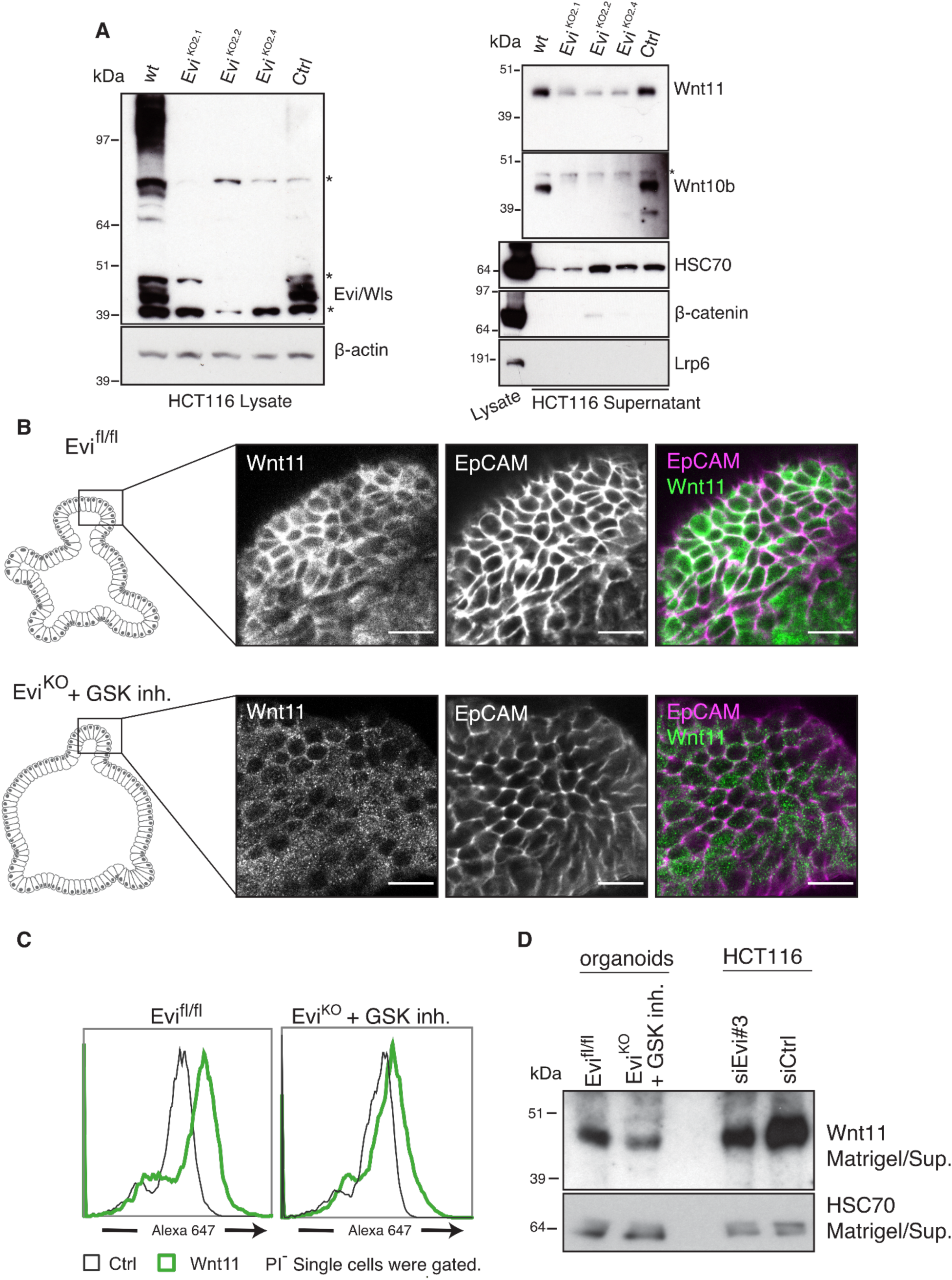
Wnt11 localizes to the membrane of Evi^KO^ intestinal organoids. **A)** Endogenous Wnt11 is still secreted by Evi^KO^ HCT116 colon cancer cells, while secretion of Wnt10b is blocked. Ctrl is a clone with wild-type Evi/Wls expression. After 72 hrs of culturing, medium was collected and subjected to Blue sepharose precipitation. Unspecific bands are marked with *. **B)** Wnt11 is present on the cell surface of wild type as well as Evi^KO^ organoids. Non-permeabilized organoids were stained with the Wnt11 antibody followed by the secondary anti-rabbit AlexaFluor488 and EpCAM-APC for membrane staining. Evi^KO^ organoids were treated with GSK3β inhibitor XVI (CHIR99021, 5 μM) to prevent their death. Scale bar is 10 µm. **C)** Wnt11 is located at the cell surface of Evi^KO^ intestinal organoids. Organoids were digested to obtain single cell suspension and stained with the Wnt11 antibody followed by the secondary antibody anti-rabbit AlexaFluor647 and propidium iodide (PI)− for detection of live cells. **D)** Wnt11 is secreted by wild type and Evi^KO^ organoids. Five day old matrigel from organoids was digested and used for Blue sepharose pull down. Samples from the supernatant of HCT116 cells transfected with the indicated siRNAs were used as controls for Wnt11 detection. HSC70 was used as the loading control. (A,D) Representative images of 3 independent experiments are shown.

Next, we asked whether Wnt11 is also secreted in the absence of the Wnt cargo receptor in an advanced primary 3D cell culture model. Therefore, we investigated Wnt11 secretion in mouse intestinal organoids. In line with our previous observations in the human colon (Voloshanenko *et al*, 2013), Evi/Wls expression in the mouse intestine is primarily found in epithelial cells located close to the bottom of the crypt and in Paneth cells (Figure S2C). In contrast, Wnt11 is highly expressed throughout the colon epithelium (Figure S2D). We generated Evi^KO^ and control intestinal organoids from Evi^fl/fl^ floxed mice (Figure S2E,F, see materials and methods) (Augustin *et al*, 2013). *In vitro* growth of mouse intestinal organoids requires Wnt signaling activity (Farin *et al*, 2012), which is impaired upon knock-out of Evi/Wls. We therefore cultured organoids in the presence of the GSK3β inhibitor CHIR99021, which activates Wnt signaling independent of Wnt secretion. To investigate whether Wnt11 is secreted in Evi^KO^ organoids, we selectively stained for Wnt11 protein in the absence of cell permeabilization, thereby only detecting protein on the cell surface (Strigini & Cohen, 2000; Farin *et al*, 2016). Extracellular Wnt11 was detected in both wild type and Evi^KO^ organoids, similar to EpCAM controls (Figure 2B). In addition, FACS analysis showed that in intact (PI-negative (PI^-^)) wild type and Evi^KO^ cells, Wnt11 was present on the cell surface (Figure 2C). Moreover, Wnt11 could be detected in the matrigel growth medium from wild type and Evi^KO^ organoids, demonstrating that the protein remains secreted under these conditions (Figure 2D). Similar to HEK293T cells, Wnt11 levels on the cell surface and cell supernatant were reduced in Evi^KO^ intestinal organoids compared to wild type cells. In summary, these experiments reveal that a substantial fraction of Wnt11 is secreted in the absence of Evi/Wls in different cell types from two mammalian species and is not simply a consequence of protein overexpression.

### Wnt11 secreted independently of Evi/Wls inhibits canonical Wnt signaling *via* FZD6

Our findings indicate that while Wnt11 and Wnt5a differ in their dependence on Evi/Wls for secretion, they have a similar ability to block Wnt3a-induced signaling in receiving cells. We therefore explored whether this inhibition is mediated by the same mechanism (Figure 3A). First, we analyzed at what level of β-catenin-dependent signaling Wnt11 interferes with Wnt3a induced signal transduction. We expressed Wnt11, treated cells with recombinant Wnt3a and analyzed activation levels of the Wnt co-receptor LRP6 and β-catenin by Western blotting. Expression of Wnt11 reduced levels of active (non-phosphorylated) β-catenin and phosphorylated LRP6, indicating that Wnt11 inhibits canonical Wnt signaling on the level of the receptor complex (Figure 3B). This effect was independent of Evi/Wls, confirming once more that Wnt11 protein secreted in the absence of the cargo receptor has signaling activity (Figure 3B).

**Figure 3.**
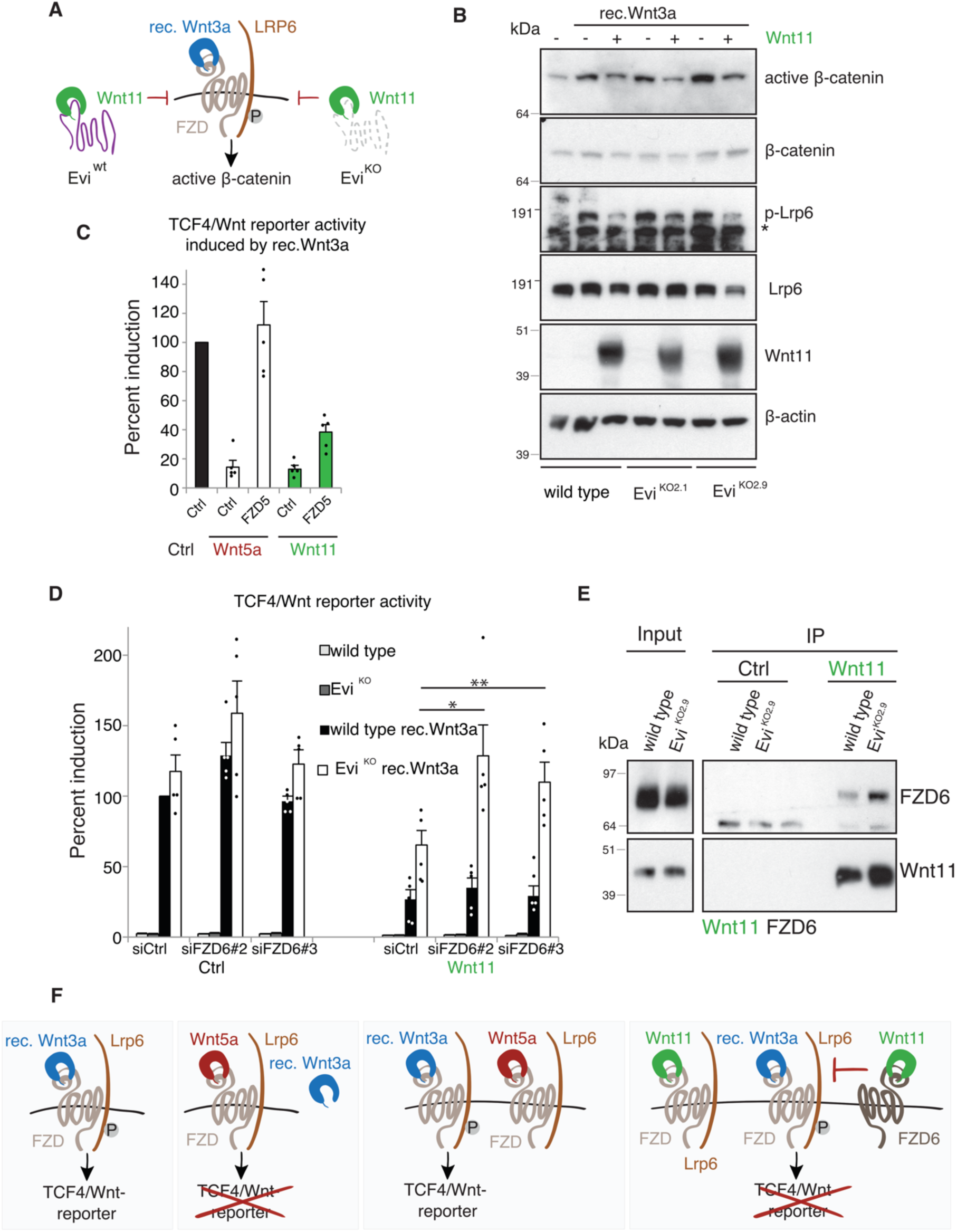
Wnt11 secreted from wild type and Evi^KO^ cells inhibits β-catenin-dependent Wnt signaling at the receptor level. **A,B**) Wnt11 inhibits β-catenin-dependent Wnt3a signaling at the receptor level. Indicated HEK293T cells were transfected with Wnt11 and lysed 48 hrs later. Recombinant mouseWnt3a (100 ng/ml) was added 16 hrs before lysis. β-actin served as the loading control. The unspecific band is marked with *. **C)** Overexpression of FZD5 rescues Wnt5a-induced inhibition of TCF4/Wnt-reporter activity, while Wnt11-induced inhibition of the reporter was just partly affected. **D)** Silencing of FZD6 diminishes the effect of Wnt11 secreted from Evi^KO^ cells. Indicated cells were reverse transfected with FZD6 siRNAs. Unpaired *t* test with Welch’s correction was performed: \**p*=0.04, \*\**p*=0.036. C,D) Indicated cells were transfected with Wnt11/Wnt5a/Ctrl and FZD5/Ctrl in (C) or Wnt11/pcDNA3 in (D) together with TCF4/Wnt-firefly luciferase and actin-*Renilla* reporters for 48 hrs. 16 hrs before the read-out recombinant mouseWnt3a (100 ng/ml) was added. Results of 5 independent experiments are shown as mean + SEM. One dot represents a single experiment. **E)** Wnt11 interacts with FZD6. HEK293T wild type or Evi^KO^ cells were stably transduced with Wnt11. These cells were transfected with the FZD6 plasmid. 48 hrs after transfection cells were lysed and Wnt11 was immunoprecipitated. (B,E) Representative results of 3 independent experiments are shown. **F)** Schematic representation of Wnt11 interactions with FZDs and FZD6 and its influence on TCF4/Wnt-reporter.

We hypothesized that the inhibitory effect of non-canonical Wnt ligands on β-catenin signaling might reflect competition of the different Wnts for Frizzled receptors. If so, then increasing receptor availability through overexpression would alleviate the inhibition. We therefore expressed different FZDs (FZD1, FZD2, FZD4, FZD5, FZD6, FZD7 and FZD10) one by one in wild type or FZD1,2,7^KO^ HEK293T cells to exclude an influence of endogenous FZDs (Voloshanenko *et al*, 2017) in combination with Wnt5a or Wnt11 and measured canonical Wnt activity (Figure 3C,S3A). Overexpression of FZD1, FZD2, or FZD5 rescued TCF4/Wnt-reporter activity when Wnt5a expressing cells were stimulated with recombinant Wnt3a, suggesting that the inhibitory effect of this Wnt is mediated by competition for receptors (Figure 3C,S3A). In contrast, inhibition of Wnt3a signaling by Wnt11 was only partially rescued by overexpression of FZD1, FZD2, or FZD5 (Figure 3C,S3A). Under these conditions, we observed still an inhibition of canonical signaling of ∼60%, suggesting that there are two pools of Wnt11, one that competes with Wnt3a for the same receptors and one that inhibits signaling through a different mechanism.

In order to identify receptors that mediate the inhibitory effect of Wnt11 that is not based on receptor competition, we silenced known receptors for non-canonical Wnts, including RoR2, PTK7, RYK, FZD6, FZD3 and CD146, and quantified TCF4/Wnt reporter levels (Figure S3D). This revealed two siRNAs targeting *FZD6* which could restore Wnt3a-induced signaling (Figure 3D). Intriguing, knockdown of FZD6 only rescued canonical Wnt signaling in Evi^KO^ cells and had no effect in wild type cells, suggesting that the Wnt11 protein secreted from cells lacking Evi/Wls signals *via* this receptor. The efficiency of FZD6 siRNAs mediating this effect directly correlated with their potency of mediating a reduction in FZD6 mRNA levels (Figure S3B).

Further analysis confirmed that the Wnt11-induced suppression of reporter activation in Evi^KO^ cells requires *FZD6*, as Wnt11 no longer suppressed the induction of the TCF4/Wnt reporter by Wnt3a when *FZD6* expression was knocked down by RNAi (Figure 3D). Silencing of *FZD6* had only a small effect on Wnt3a-induced TCF4/Wnt reporter activity in the absence of Wnt11 both in wild type and Evi^KO^ cells (Figure 3D). Furthermore, the effect of Wnt11-mediated inhibition on canonical Wnt signaling was enhanced upon overexpression of FZD6 in Evi^KO^ cells, which stably express Wnt11 (Figure S3C), and a partial rescue was observed when overexpressing FZD6 in the presence of siFZD6#3 in these cells (Figure S3C).

We also tested if Wnt11 binds to FZD6 by co-immunoprecipitation experiments. Overexpressed FZD6 was pulled down with Wnt11 in wild type and Evi^KO^ cells, with the interaction particularly prominent in the absence of the cargo receptor (Figure 3D). Together, these experiments indicate that the Evi/Wls-dependent Wnt11 fraction is competing with Wnt3a for binding to FZD receptors, thereby inhibiting its signaling activity. In contrast, Wnt11 that is secreted independent of Evi/Wls binds to FZD6 and inhibits canonical signaling through this receptor (Figure 3F).

### Wnt11 secreted from Evi^KO^ is not lipidated by C16:1 palmitoleic acid

Our finding that Wnt11 is secreted in an Evi/Wls dependent and an Evi/Wls independent fraction suggests that this Wnt protein might partially evade binding to its cargo receptor. To examine potential differences in Evi/Wls binding between Wnt11 and Wnt5a, whose secretion strictly depends on the Evi/Wls, we performed V5 immunoprecipitation using overexpressed V5 tagged proteins in HEK293T cells. While both Wnts bound to Evi/Wls, Wnt11 showed strongly reduced binding compared to Wnt5a, despite comparable expression levels of both proteins (Figure 4A).

**Figure 4.**
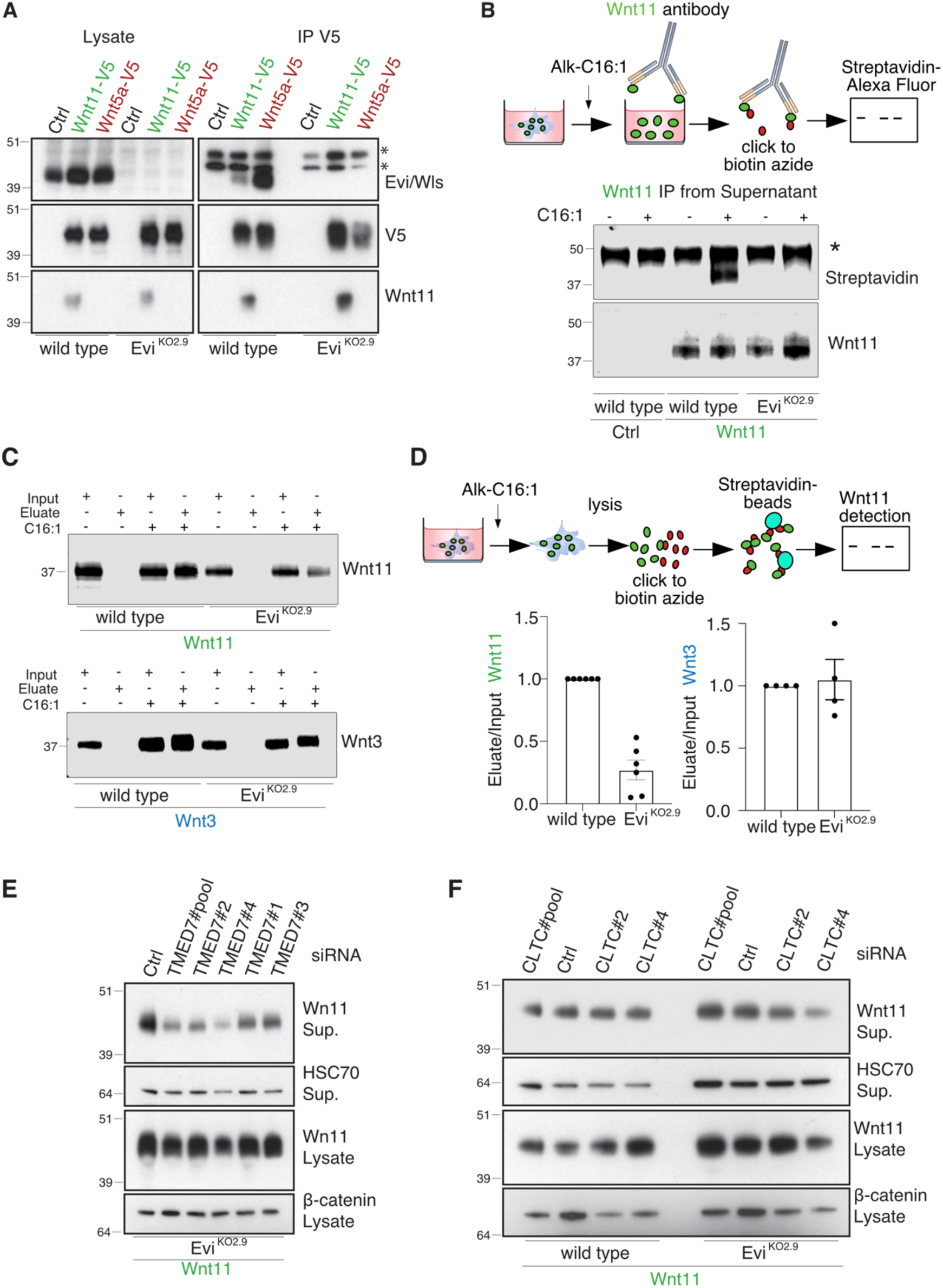
Wnt11 secreted by Evi^KO^ is not palmitoleoylated. **A)** Wnt5a binds Evi/Wls stronger than Wnt11. Indicated HEK293T cells were transfected with Wnt5a-V5 or Wnt11-V5. 48 hrs after transfection, Wnts were immunoprecipitated by V5-agarose. **B)** Wnt11 secreted from wild type cells is C16:1 lipidated, while palmitoleoylation is not detected in the case of Wnt11 secreted from Evi^KO^ cells. Indicated cells were transfected with Wnt11 and pre-incubated with Alk-C16:1 for 24 hrs. Then Wnt11 was immunoprecipitated from the supernatant and the click reaction was performed on the beads. Presence of Wnt11 palmitoleoylation was detected by streptavidin. * unspecific binding of streptavidin. Representative results of 6 independent experiments are shown. **C,D)** Evi^KO^ cells have less palmitoleoylated Wnt11 than wild type cells. Indicated HEK293T cells were transfected with Wnt11 or Wnt3. 24 hrs post-transfection, cells were pretreated with Alk-C16:1 for 24 hrs. The click reaction using biotin azide was performed in cell lysates followed by the enrichment of lipidated proteins by streptavidin beads. Input and eluate from the beads were loaded and the presence of Wnt11 or Wnt3 was detected by WB. D) Quantification of the ratio between eluate and input are shown on the graphs. Results of 6(Wnt11)/4(Wnt3) independent experiments are shown as mean +- SEM. One dot represents a single experiment. **E,F)** Silencing of TMED7 but not CLTC leads to the reduction of secretion of Wnt11. Cells were transfected with the indicated siRNAs for 72 hrs and with Wnt11 for 48 hrs. Wnt11 was precipitated from supernatant (Sup.) with Blue sepharose. β-catenin served as the loading control in the case of the lysates, while HSC70 was used as the loading control for the supernatant. E,F) Representative results of 3 independent experiments are shown.

Current evidence suggests that Wnt proteins rely on Evi/Wls for cellular export due to their lipid modification, with structural data showing extensive interactions between the lipid moiety of Wnts and the cargo receptor (Zhong *et al*, 2021; Nygaard *et al*, 2021). We therefore investigated whether the Wnt11 protein secreted in the absence of Evi/Wls is modified by C16:1 palmitoleic acid (Alk-C16:1). To test the palmitoleoylation status of Wnt11 in the cell supernatant, cells were metabolically labeled with Alk-C16:1. Afterward, Wnt11 from the supernatant of wild type and Evi^KO^ HEK293T cells was immunoprecipitated and biotin was added to the lipid chain *via* a click reaction. Detection of biotinylated protein revealed that Wnt11 secreted by wild type cells was palmitoleoylated, while Wnt11 secreted from HEK293T Evi^KO^ cells was not (Figure 4B). Next, we checked the palmitoleoylation of Wnt11 in whole cell lysates. HEK293T wild type and Evi^KO^ cells were labeled with Alk-C16:1. Click to biotin was performed in cell lysates followed by enrichment of lipidated proteins *via* streptavidin and detection of Wnt proteins in Western blot analysis. This revealed that palmitoleoylation of Wnt11 is reduced in the absence of Evi/Wls, while such a reduction was not detected for the canonical Wnt3 protein (Figure 4C). Together, these data suggest that a fraction of Wnt11 is not palmitoleoylated, providing a mechanistic explanation for the observed reduced binding to Evi/Wls and the ability to be secreted in its absence.

Next, we investigated whether in the absence of Evi/Wls secretion of Wnt11 depends on other proteins. p24 proteins are cargo receptors that cycle between the ER and Golgi and have been implicated in Wnt export (Port *et al*, 2011; Buechling *et al*, 2011; Li *et al*, 2015). Interestingly, *Drosophila* WntD (Gordon *et al*, 2005), which can be secreted without Evi/Wls (Ching *et al*, 2008), requires the p24-protein Opossum for cellular export (Buechling *et al*, 2011). Consistent with these findings, we found that knockdown of the human p24 family member TMED7 (p27) results in a marked reduction of Wnt11 secretion in Evi^KO^ cells (Figure 4E). We also investigated the importance of clathrin-mediated endocytosis, as this step has been implicated in Wnt secretion in various contexts (Port *et al*, 2008; Pan *et al*, 2008). Knock-down of clathrin heavy chain (CLTC) with several independent siRNAs (Figure S4A) strongly reduced Wnt3 secretion (Figure S4B), while secretion of Wnt11 was not affected in wildtype or Evi^KO^ cells (Figure 4E), indicating once more that these proteins are secreted through different routes.

### The N-terminus of Wnt11 mediates Evi/Wls-independent secretion

To gain insights into the molecular determinants that mediate the differential dependency of different Wnt ligands on Evi/Wls, we generated chimeric constructs of Wnt11 and Wnt3 (Figure 5A). Previous work showed that the N-terminus of non-canonical Wnts (Wnt5a and Wnt11) is responsible for the activation of different branches of β-catenin-independent signaling in Wnt receiving cells in *Xenopus* (Wallkamm *et al*, 2016). Therefore, we hypothesized that this part of the protein might also explain the observed differences in secretion of Wnt3 and Wnt11. We first tested a chimeric Wnt3^1-115^-Wnt11 protein (Figure 5A,S5A), consisting of the first 115 amino acids of Wnt3 followed by 240 aa of Wnt11. Secretion of Wnt3^1-115^-Wnt11 was strongly reduced in Evi^KO^ cells compared to wild type cells (Figure 5B), resembling the behavior of Wnt3. This suggests that the N-terminus of Wnt proteins determines their dependence on the Evi/Wls cargo receptor.

**Figure 5.**
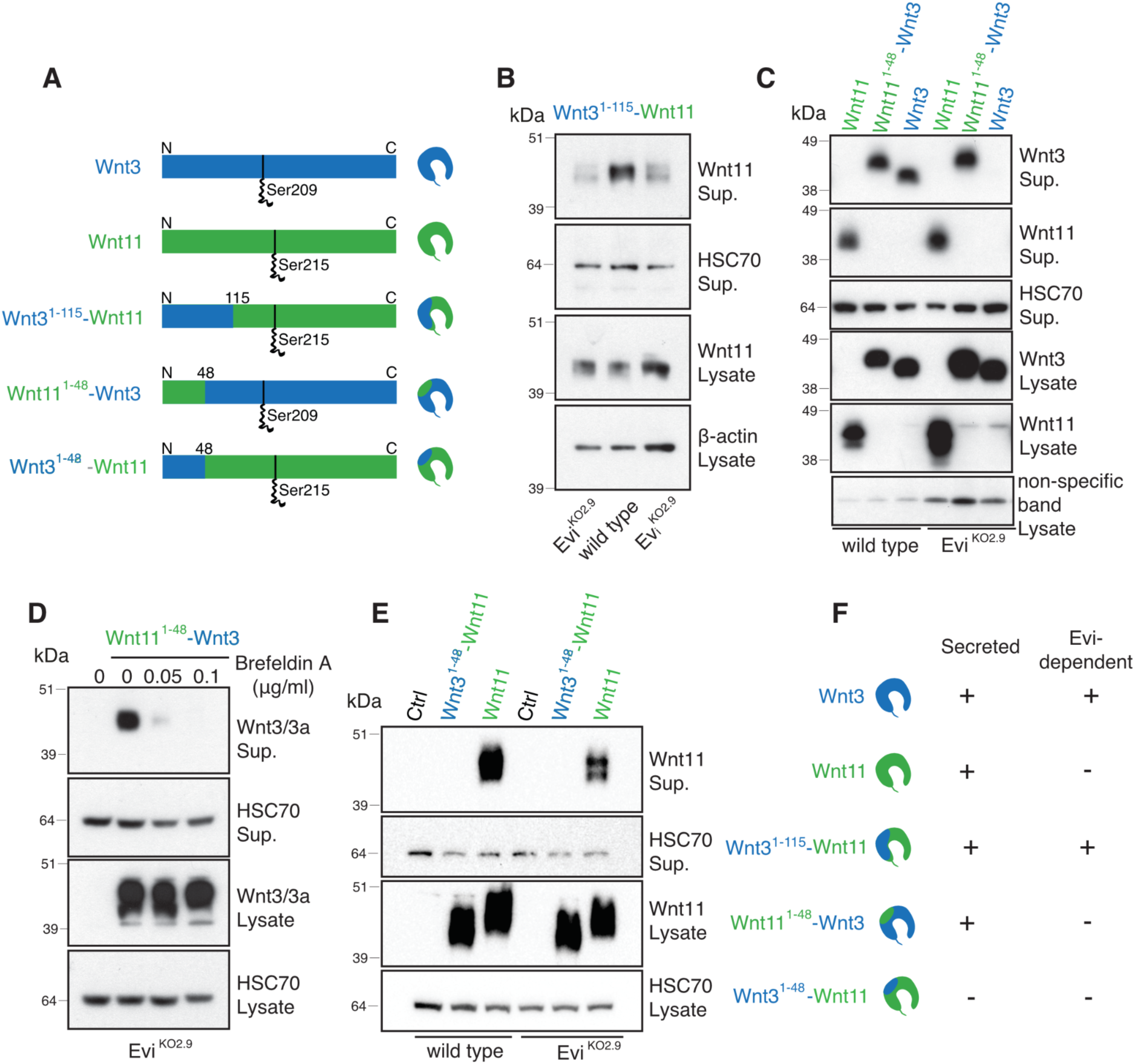
N-terminus of Wnt11 determines Evi/Wls-independent secretion. **A)** Schematic representation of Wnt11, Wnt3 and their chimeras. **B,C)** Secretion of Wnt3^1-115^-Wnt11 chimeric protein is blocked in Evi^KO^ cells, while Wnt11^1-48^-Wnt3 chimera is secreted. **D)** Wnt11^1-48^-Wnt3 secretion is blocked by Brefeldin A. Evi^KO^ HEK293T cells were transfected with Wnt11^1-48^-Wnt3 for 24 hrs and then cells were treated with indicated amounts of Brefeldin A for 20 hrs. **E)** Wnt3^1-48^-Wnt11 is expressed but not secreted. B-E) Indicated HEK293T cells were transfected with the Wnt3^1-115^-Wnt11, Wnt11^1-48^-Wnt3 or Wnt3^1-48^-Wnt11 constructs. After 48 hrs supernatants (Sup.) were subjected to Blue Sepharose pulldown. HSC70 served as the loading control. Representative WBs of 3 independent experiments are shown. **F)** Schematic representation of Wnt3, Wnt11 and chimeric proteins and their dependency on Evi/Wls.

To gain further insights into the underlying mechanism, we compared the first 115 aa of Wnt11 to those of other Wnt ligands. This revealed one region (QECQHQFRG in Wnt3), which was highly homologous between different Wnts, but divergent in Wnt11 (Figure S5B). However, a Wnt11 variant with this motif replaced with the amino acids present in Wnt3 was still secreted in the absence of Evi/Wls (Figure S5C). We then focused on the N-terminal sequence of Wnt11, which is the other region of the protein that is divergent from other Wnts (Figure S5B). We generated a chimeric protein, Wnt11^1-48^-Wnt3, in which the first 48 N-terminal amino acids of Wnt11 replace the respective amino acids of Wnt3 (Figure S5A,5A). Interestingly, Wnt11^1-48^-Wnt3 was efficiently secreted from Evi^KO^ cells, phenocopying Wnt11 rather than Wnt3 (Figure 5C), a process that could be inhibited by Brefeldin A (Figure 5D). We also generated the complementary chimeric protein, Wnt3^1-48^-Wnt11, which was expressed, but not secreted, by both wild type and Evi^KO^ cells, suggesting general problems with protein trafficking or folding (Figure 5E). Together, our data suggest that the first 48 amino acids of Wnt3 and Wnt11 are the key determinants for the dependence on Evi/Wls for secretion (Figure 5F).

### Asparagine 40 of Wnt11 reduces its binding affinity for Evi/Wls

Our data indicate that the ability of Wnts to be secreted with or without Evi/Wls is reflected in the affinity of the ligands for the cargo receptor (Figure 4A). We therefore tested our chimeric Wnt proteins for Evi/Wls binding. Co-immunoprecipitation analysis revealed that wild type Wnt3 bound Evi/Wls stronger than Wnt11^1-48^-Wnt3 (Figure 6A). Conversely, Wnt3^1-48^-Wnt11, while not secreted, was able to precipitate much more Evi/Wls than Wnt11 (Figure 6B). These data indicate that the N-terminus of Wnts strongly modulates the binding affinity for Evi/Wls.

**Figure 6.**
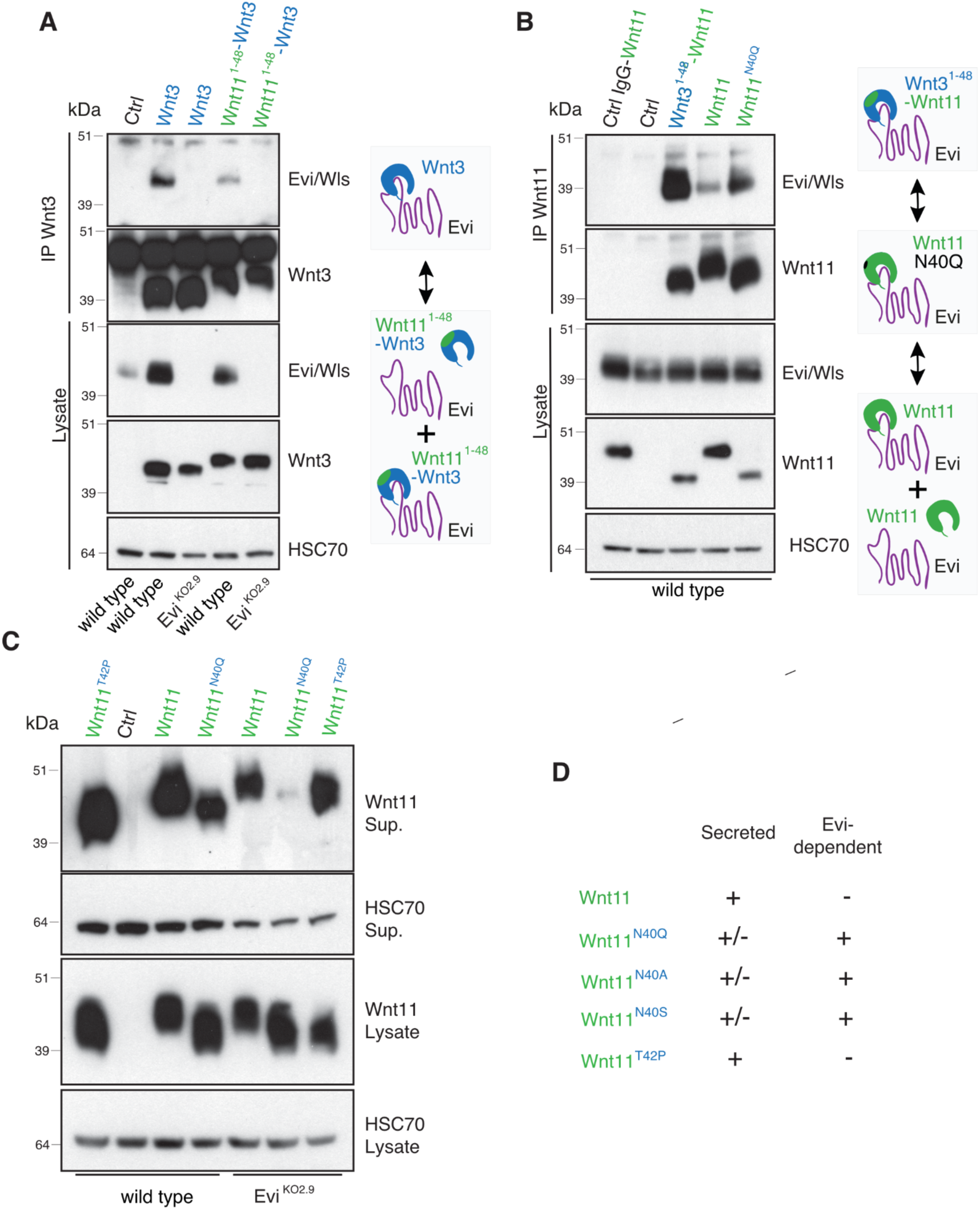
Wnt-binding affinity for Evi/Wls is the crucial regulator for Evi/Wls-independent secretion. **A)** Wnt3 has stronger affinity for Evi/Wls than Wnt11^1-48^-Wnt3. **B)** Wnt3^1-48^-Wnt11 and Wnt11^N40Q^ bind Evi/Wls stronger than Wnt11. **C)** Wnt11^N40Q^ is not secreted by Evi^KO^ cells. Wnts were precipitated from supernatant (Sup.) with Blue sepharose. A-C) HEK293T wild type or Evi^KO^ cells were transfected with the indicated plasmids for 48 hrs. HSC70 served as the loading control. Representative results of 3 independent experiments are shown. **D)** Secretion and dependency on Evi/Wls of indicated proteins.

Interestingly, we observed that the migration behavior in gel electrophoresis of the chimeric proteins Wnt11^1-48^-Wnt3 and Wnt3^1-48^-Wnt11 was different compared to Wnt3 and Wnt11 (Figure 6A,B). While Wnt11^1-48^-Wnt3 runs above Wnt3 on the gel, Wnt3^1-48^-Wnt11 runs below Wnt11. This difference could not be simply explained by mass changes due to the different amino acid composition, so we surmised that there might be differences in post-translational modifications. Previously, it has been shown that glycosylation of Wnt3a is required for subsequent palmitoleoylation and secretion (Komekado *et al*, 2007). To investigate whether glycosylation might explain the observed migration pattern, we treated lysates from Wnt3, Wnt11^1-48^-Wnt3, Wnt11 and Wnt3^1-48^-Wnt11 expressing cells with a mix of deglycosylation enzymes. Deglycosylation decreased migration of Wnt3, Wnt11^1-48^-Wnt3, Wnt11 and Wnt3^1-48^-Wnt11 to the same level as non-treated controls (Figure S6A).

It has been previously described that Wnt11 is glycosylated at Asparagine 40 (Yamamoto *et al*, 2013). To test whether this residue is responsible for the observed differences in migration and secretion, we mutated asparagine 40 to glutamine. Indeed, Wnt11^N40Q^ migrated similar to Wnt3^1-48^-Wnt11 on the gel, indicating this residue is responsible for the differential glycosylation. Importantly, secretion of Wnt11^N40Q^ in wild type cells was reduced compared to Wnt11 and blocked in Evi^KO^ cells (Figure 6C). These results suggested that the glycosylation of asparagine 40 in Wnt11 leads to reduced Evi/Wls affinity and secretion of a substantial fraction of the protein in the absence of the cargo receptor. Alternatively, the amino acid itself could have a structural role independent of its post-translational modification. To distinguish between these possibilities, we generated a Wnt11^T42P^ mutant, as asparagine-linked glycosylation typically requires the consensus sequences (Asn-X-Ser/Thr) (Breitling & Aebi, 2013). As expected, Wnt11^T24P^ showed similar changes in migration, consistent with reduced glycosylation. Surprisingly, the mutant was still secreted by Evi^KO^ cells at similar levels than wild type Wnt11, suggesting that asparagine 40, and not its glycosylation, is the key determinant of Evi/Wls independent secretion (Figure 6C). To confirm this, we mutated the asparagine in position 40 to either alanine or serine, which is present at this position in Wnt3. Secretion of both Wnt11^N40S^ and Wnt11^N40A^ mutants was inhibited in Evi^KO^ cells, similar to Wnt11^N40Q^ (Figure S6B). We also generated a double mutant, Wnt11^N40Q,T42P^, which showed a migration behavior similar to Wnt11^N40Q^ and Wnt11^T42P^ (Figure S6B). As Wnt3^1-48^-Wnt11 binds more Evi/Wls than Wnt11, we also analyzed the interaction of Evi/Wls with Wnt11^N40Q^. Similar to Wnt3^1-48^-Wnt11, more Evi/Wls was detected upon immunoprecipitation of Wnt11^N40Q^ than compared to Wnt11 (Figure 6B).

Together, this identifies asparagine 40 as a crucial amino acid modulating binding affinity of Wnt11 for Evi/Wls. While most Wnts have high binding affinity for Evi/Wls and are immediately captured after synthesis, a fraction of Wnt11 has lower binding affinity and is able to escape Evi/Wls and be secreted without it.

### The Wnt11 N-terminus can switch Wnt3 signaling activity

The N-terminal 48 amino acids of Wnt11 are sufficient to alter the dependency of Wnt3 on Evi/Wls for secretion. We therefore asked if such a chimeric protein would also resemble Wnt11 when inducing downstream Wnt signaling in receiving cells. First, we tested this protein in the TCF4/Wnt-reporter assay, which is robustly activated by Wnt3, but not Wnt11. Wnt11^1-48^-Wnt3 did not induce any activation of the canonical TCF4/Wnt-reporter (Figure 7A). Next, we asked whether this chimeric protein can inhibit β-catenin-induced signaling similar to Wnt11. Indeed, expression of Wnt11^1-48^-Wnt3 inhibited recombinant Wnt3a-induced TCF4/Wnt reporter activity in wild type and in Evi^KO^ cells with the same efficiency as Wnt11 (Figure 7B). Furthermore, overexpression of classical FZDs (FZD2 and FZD5) was not able to rescue this inhibition of reporter activity (Figure 7C,S7A). Together, these experiments show that in HEK293T cells Wnt11^1-48^-Wnt3 has the same signaling properties as Wnt11, suggesting that the N-terminus of the protein has a strong impact on receptor binding or activation.

**Figure 7.**
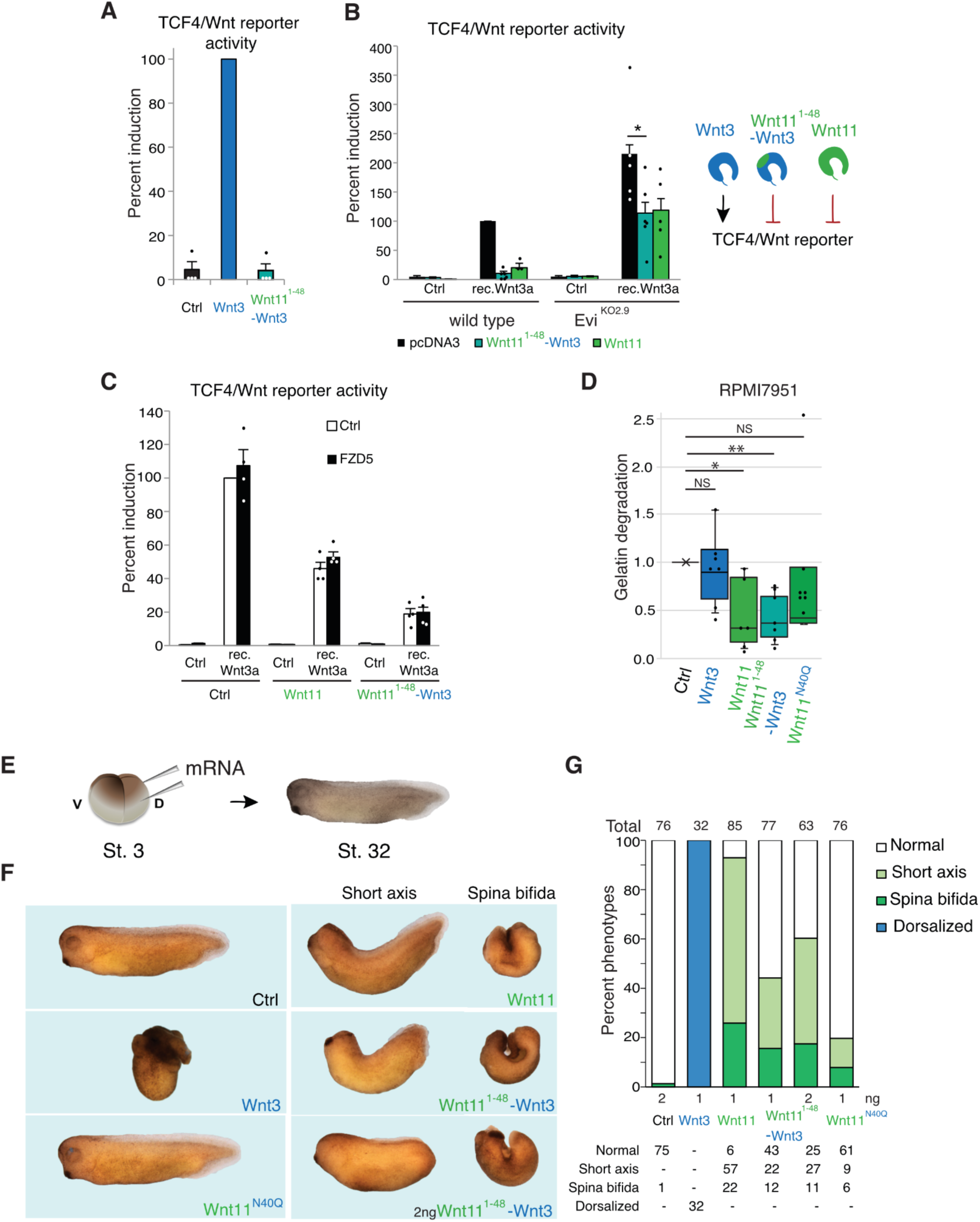
N-terminus of Wnt11 switches Wnt3 function from canonical to non-canonical. A,B) Wnt11^1-48^-Wnt3 chimera inhibits Wnt3a-induced TCF4/Wnt-reporter activity. **C)** FZD5 overexpression does not rescue Wnt11^1-48^-Wnt3-induced inhibition of rec. Wnt3a-induced TCF4/Wnt-reporter activity. A-C) HEK293T were transfected with the indicated pcDNA3, Wnt3, FZD5, Wnt11^1-48^-Wnt3 or Wnt11 constructs together with TCF4/Wnt-firefly luciferase and actin-Renilla reporters for 48 hrs. 100 ng/ml recombinant mouse Wnt3a was added 16 hrs before the read-out. Results of 4(A,C) or 5/6(B) independent experiments are shown as mean + SEM. One dot represents a single experiment. * *p* =0.0459 (Unpaired *t* test with Welch’s correction). **D)** Wnt11 and Wnt11^1-48^-Wnt3 reduce gelatin degradation capacity of RPMI-7951 melanoma cells. Cells were transfected with Ctrl, Wnt3, Wnt11, Wnt11^1-48^-Wnt3 or Wnt11^N40Q^ for 72 hrs and seeded on fluorescein-gelatin (green) coated coverslips. After 24 hrs, cells were fixed and stained for DNA (blue) and actin (orange). Box plots show the averaged “gelatin degradation capacity per cell” normalized to its control treatment in at least 6 independent replicates with a minimum of 100 cells per condition. One-Sample Wilcoxon Signed Rank Test, * *p* =0.031, ** *p* =0.0156, NS-not significant. **E,F)** The N-terminus of Wnt11 induces a non-canonical mode of action for Wnt3 in *X. laevis.* E) Microinjection strategy and F) representative phenotypes of *X. laevis* tadpoles at NF stage 32. Embryos were classified as “spina bifida”, “short axis” or “normal” according to their phenotypical characteristics. **G)** Quantification of 3 independent injections.

Next, we investigated the signaling properties of Wnt11^1-48^-Wnt3 in a different cellular context. In melanoma, both canonical and non-canonical Wnt signaling contribute to the onset and progression of tumorigenesis in a complementary and opposing manner (Webster *et al*, 2015; O’Connell & Weeraratna, 2009). Previous studies showed that the non-canonical Wnt5a promotes invasion and metastasis in melanoma cells (Weeraratna *et al*, 2002). Invasiveness of cells *in vitro* can be visualized by seeding cells on fluorescently-labeled extracellular matrix (Lu *et al*, 2016). We overexpressed Wnt3, Wnt11, as well as Wnt11^1-48^-Wnt3 and Wnt11^N40Q^, in RPMI-7951 melanoma cells and seeded them on fluorescein-labeled gelatin. Whereas overexpression of Wnt3 or Wnt11^N40Q^ did not change gelatin degradation compared to the control, overexpression of both Wnt11 and Wnt11^1-48^-Wnt3 reduced gelatin degradation significantly (Figure 7D, S7B).

Finally, we tested the signaling activity of Wnt11^1-48^-Wnt3 *in vivo*. Previously, it has been shown that in *Xenopus* Wnt5a and Wnt11 regulate convergent extension (CE) movements (Tada & Smith, 2000; Yamanaka *et al*, 2002). To investigate the role of the N-terminus of Wnt11 in this context, mRNAs of Wnt3, Wnt11, Wnt11^1-48^-Wnt3 and Wnt11^N40Q^ were injected into *Xenopus* embryos. As expected, injection of Wnt3 mRNA caused severe dorsalization, while injection of Wnt11 mRNA resulted in a high incidence of tadpoles with a shortened axis or spina bifida (Fig. 7E-G). Injection of *Wnt11^1-48^-Wnt3* mRNA also resulted in a shorter axis and *spina bifida*, hallmarks of misregulated CE movements, and similar to embryos injected with *Wnt11* mRNA. This effect was concentration-dependent, as increasing the amount of mRNAs increased the effect (Figure 7G). Furthermore, injection of Wnt11^N40Q^ mRNA caused a less severe phenotypes than Wnt11 mRNA, confirming the critical role of this amino acid in Wnt11 activity (Figure 7G). To assess Wnt/PCP signaling activity in *Xenopus* embryos, we used an ATF2 luciferase reporter assay (Figure S7C)(Ohkawara & Niehrs, 2011). Both *Wnt11^1-48^-Wnt3* and *Wnt11* mRNAs increased Wnt/PCP signaling, while *Wnt3* and *Wnt11^N40Q^* mRNAs had no effect.

In summary, our data show that the first 48 amino acids of Wnt11 are sufficient to render Wnt3 partially independent of the cargo receptor Evi/Wls, but also profoundly alter its signaling properties. We find that in various *in vitro* and *in vivo* assays, a *Wnt11^1-48^-Wnt3* hybrid protein phenocopies Wnt11, suggesting that this portion of the protein dictates whether a Wnt signals *via* the canonical or non-canonical signaling pathway.

## DISCUSSION

In this study, we demonstrate that Wnt11 can be secreted independently of Evi/Wls, a pan-Wnt cargo transporter thought to be generally required for secretion of palmitoleoylated Wnt proteins. We found that the ability of Wnt11 to be secreted in the absence of Evi/Wls is encoded in its N-terminus. Strikingly, the N-terminal region of Wnt11 is also sufficient to strongly modulate the interaction of other Wnt proteins with the cargo receptor and influences their signaling to downstream cells. A chimeric Wnt11^Nterm^-Wnt3 protein shows reduced binding affinity for Evi/Wls and switches the canonical Wnt protein to induce non-canonical signaling. Moreover, during *Xenopus laevis* development, the chimeric Wnt11^Nterm^-Wnt3 protein phenocopies Wnt11 and induces defects in convergent extension movement.

The present scientific view is that Wnt proteins are lipid modified glycoproteins that rely on Evi/Wls for their secretion. Until recently, the only known exception was *Drosophila* WntD, which lacks the serine-residue required for lipid-modification, is not glycosylated and is secreted independently of Evi/Wls (Ching *et al*, 2008). The lack of lipid modification might also underlie WntD’s unique signaling properties. WntD does not regulate the canonical Wnt pathway, but instead inhibits Dorsal/NF-kB signaling (Ganguly *et al*, 2005; Gordon *et al*, 2005) and the Toll pathway *via* its interaction with FZD4 (Rahimi *et al*, 2016). Recently, it was shown that *Xenopus* Wnt8 can be secreted without Porcupine and retains its activity upon high expression levels (Speer *et al*, 2019). In addition, a fraction of Wnt7a can be secreted on extracellular vesicles in a process that is not entirely dependent on Evi/Wls (Gurriaran-Rodriguez *et al*, 2024). In line with these studies, we show here that mammalian Wnt11 is also in part not lipid modified and secreted independently of Evi/Wls. Together, a picture emerges that Wnt protein secretion is more complex than initially anticipated, with some Wnts being secreted by multiple routes.

One of the fundamental questions in Wnt protein research is identifying the structural determinants that underlie their different biological activities. Our data demonstrate that Evi/Wls-independent secretion is more widespread than anticipated, but not a general property of Wnt ligands that elicit non-canonical signaling, as Wnt5a secretion is strictly Evi/Wls-dependent. Instead, our results indicate that the amino acid composition of the Wnt N-terminus plays an important role in determining cargo receptor binding. Recent cryo-EM structures of Evi/Wls in complex with Wnt proteins have revealed the presence of specific hairpins in Wnts, which form critical interactions with the carrier (Zhong *et al*, 2021; Qi *et al*, 2023; Nygaard *et al*, 2021). The superposition of the Wnt11 AlphaFold2 (AF2) (Jumper *et al*, 2021) model onto Wnt3a (PDB ID 7DRT) shows a similar fold (Figure S8A). Nevertheless, the differences in the primary sequences of the Wnt3/Wnt11 N-termini might also influence post-translational modifications essential for PORCN and Evi/Wls binding. In an AF2 multimer prediction of the PORCN-Wnt11 complex, the asparagine in position 40 (N40) of Wnt11 is in proximity to the PORCN region N222-A234 (Figure S8B), suggesting this interaction could modulate Wnt11 binding to PORCN. It is therefore tempting to speculate that the PORCN-Wnt11 interaction might affect both Wnt lipidation efficiency and subsequent Evi/Wls loading. Alternatively, N40 might directly modulate Evi/Wls affinity of Wnt11. In an AF2 multimer prediction of the Evi/Wls-Wnt11 complex, N40 is about 9 Å apart from the Wnt-binding domain (Figure S8C).

Wnt ligands induce diverse signaling responses in signal receiving cells, but how differences in Wnt protein composition are translated into differences in downstream signaling remains poorly understood. Sequence alignment of different Wnt ligands showed that despite overall high homology, the N-terminal sequence is divergent between Wnt ligands (Figure S5B). However, whether this correlates with the function of the different Wnts is unclear. Our data show that the N-terminus of Wnt11 can play an instructive role for signaling specificity when transplanted into a different Wnt ligand. In line with our data, in *Xenopus* it has been shown that the N-terminal part of Wnts is essential for non-canonical Wnt signaling and the C-terminus for canonical Wnt activity (Wallkamm *et al*, 2016). Interestingly, alternative splicing events might create different N-termini for a variety of Wnt ligands, like Wnt16 and Wnt5a (Schubert *et al*, 2025; Katoh & Katoh, 2005). Therefore, alternative splicing might result in isoforms that can either induce canonical or non-canonical Wnt signaling.

Canonical Wnt signaling is characterized by translocation of β-catenin to the nucleus and subsequent transcriptional regulation, while non-canonical pathways function *via* diverse mechanisms. A common hallmark of non-canonical signaling is the inhibition of β-catenin-mediated signaling. As indicated by a previous study, Wnt11 can inhibit canonical Wnt signaling through different mechanisms, including transcriptional repression and competing with canonical Wnts for FZD binding (Maye et al. 2004). Our findings suggest that these different modes of Wnt11-mediated inhibition reflect distinct pools of the protein secreted from producing cells. While lipid-modified and Evi/Wls-dependent Wnt11 competes with canonical Wnt proteins for FZD binding, non-palmitoleoylated Wnt11 secreted independently of Evi/Wls inhibits β-catenin signaling even in the presence of excess canonical Wnt receptors. This alternative inhibitory mechanism operates through FZD6 signaling. The ability to suppress canonical Wnt signaling through non-competitive mechanisms is encoded in the Wnt11 N-terminus, as demonstrated by a chimeric protein containing the Wnt11 N-terminus fused to Wnt3.

In summary, our study expands the emerging view that Wnt proteins are secreted through more diverse mechanisms than previously anticipated. Mechanistically, we demonstrate that the N-terminus of Wnts, a particularly variable region within this protein family, encodes critical information governing their lipid modification, secretion, and downstream signaling specificity. The existence of multiple secretion pools for a single Wnt ligand could provide a mechanism for fine-tuning signaling strength and duration within developmental contexts. Moreover, the ability of differentially processed Wnt proteins to engage distinct downstream pathways suggests that cells can deploy the same ligand to achieve fundamentally different biological outcomes, adding a new layer of regulatory complexity to Wnt-mediated developmental programs.

## MATERIAL AND METHODS

### Cell Culture

HEK293T and RPMI-7951 melanoma cells were cultured in Dulbecco’s MEM (GIBCO, Life Technologies GmbH, Darmstadt, Germany) supplemented with 10% fetal bovine serum (Biochrom GmbH, Berlin, Germany) without antibiotics. Colon cancer HCT116 cells were cultured in McCoy’s medium (GIBCO, Life Technologies, Darmstadt, Germany) supplemented with 10% fetal bovine serum. Cell lines were obtained from ATCC (Washington, USA) and authenticated by Multiplexion (Heidelberg, Germany). All cell lines were regularly confirmed to be mycoplasma negative.

### Short-guide (sg) RNA design and vector cloning

Short-guide RNAs for CRISPR/Cas9 experiments were designed using E-CRISP (http://www.e-crisp.org) (Heigwer *et al*, 2014). sgRNA sequences used in this study are listed in Supplementary Table S3. Oligonucleotides were purchased from Eurofins (Ebersberg, Germany), annealed and cloned into the *Bbs1*–*Bbs1 (BpiI-BpiI*, Fermentas/Thermo Fisher Scientific, Darmstadt, Germany) sites downstream from the human U6 promoter in px459 vector (Addgene #48139) according to (Ran *et al*, 2013).

### Generation of gene knockout cell lines

HEK293T and HCT116 pools were generated as previously described (Ran *et al*, 2013). In brief, cells were transfected with px459 vectors expressing sgRNA and Cas9 using TransIT-LT1 transfection reagent (731-0029; VWR, Darmstadt, Germany) according to the manufacturer’s description. Efficiency of transfection was checked by transfection of RFP expressing plasmid and counting of RFP-positive cells in independent wells. 48 hrs after transient transfection, transfected cells were selected with 2 μg/ml of puromycin (P9620, Sigma-Aldrich) for 30 hrs until all cells transfected with the control plasmid died. For experiments with pools of HEK293T, cells were cultured for 4-7 days before starting the experiment. For generation of cell clones, cells were diluted to a concentration of 500 cells per 1 ml and seeded in 220 µl medium per well in a 96-well-plate following four steps of serial dilution (1:10). Single cell clones were identified by microscopy and subsequently expanded. HCT116 Evi^KO^ clones were expanded in the presence of recombinant mouse Wnt3a (50 ng/ml, Peprotech, Hamburg, Germany) or 20% filtered medium of wild type HCT116 cells.

### Mutation analysis by indel-nested PCR

Knock-out in selected clonal cell lines was confirmed by amplicon sequencing. The 200-300 bp regions of interest were selected based on predicted sgRNA target sites and primers for amplification were generated using Primer3Web (Untergasser *et al*, 2012). For amplicon sequencing, primer adapters were added (see also Supplementary Table S5). We validated primers using DNA of wild type cells. Genomic DNA was isolated with DNeasy Blood and Tissue Kit (Qiagen, Hilden, Germany) according to the manufacturer’s instructions. Primers described in Supplementary Table S6 were used for the first amplification step (20 cycles). After purification with the PCR Cleanup Kit (Machery-Nagel, Düren, Germany) the second amplification step was performed using the PCR primers described in Supplementary Table S5. PCR products were purified as described above, and the DNA concentration was detected by Qubit dsDNA HS Assay Kit (Q32854; Life Technologies, Darmstadt, Germany). Samples were diluted, pooled and sequenced on an Illumina Miseq sequencing platform.

### Generation of mutant Wnt constructs

To generate the chimeric Wnt constructs, pcDNA Wnt11 and pcDNA Wnt3 were used as templates (Najdi *et al*, 2012). gBlocks (Supplementary Table S12) of the chimeric parts, which are different from the original vectors, were purchased from IDT (Iowa, USA) and cut with the same enzymes (Supplementary Table S12) as the corresponding vectors. gBlocks and vectors were ligated in 5:1 ratio.

For generation of point mutations, the QuikChange II Site-directed mutagenesis kit (Agilent, CA, USA) was used according to the manufacturer’s instructions. Primers are listed in Supplementary Table S11.

### Wnt secretion assay

The Wnt cDNA plasmids used in this study are listed in the Supplementary Table S4 (MacDonald *et al*, 2014; Najdi *et al*, 2012). Transfection of HEK293T cells was performed in 6 well-plates using TransIT-LT1 transfection reagent (731-0029; VWR, Darmstadt, Germany). 48h after transfection, supernatants (2 ml) were collected and debris was removed by centrifugation. Next, 1% TritonX-100 with Blue Sepharose:PBS (1:1; 15 μl per 1 ml) were added to the supernatant. Precipitation of Wnt ligands was done for 20-24 hrs at 4 °C on a rotating platform. Blue sepharose was washed (PBS + 1% TritonX-100) once for 10 min and proteins were eluted with 1x loading dye (50 μl) at 70 °C for 10-15 min with shaking. For detection of endogenous Wnts, HCT116 cells were washed 3-4 times with PBS to remove remaining Wnts in the medium and incubated with fresh medium for 72 hours. 10 ml of supernatant of HCT116 cells was used for analysis.

### Western blot analysis

Cells were collected by trypsinization and washed with PBS. Subsequently, cell pellets were lysed in 100-200 µl lysis buffer (20 mM TRIS-HCL pH7.4, 130 mM NaCl, 2 mM EDTA, 1% TritonX-100 and complete protease inhibitors (Roche, Basel, Switzerland)). For the detection of p-Lrp6 and active-β-catenin, phosphatase inhibitor cocktails 2 (P5726; Sigma, St. Louis, MO, USA) and 3 (P0044, Sigma) were added to the lysis buffer according to the manufacturer’s instructions. Cells were lysed on ice for 30 min and samples were centrifuged at 16000g at 4 °C for 30 min. After centrifugation, the supernatant was collected and a loading buffer was added. Samples were heated at 70 °C for 10 min and loaded on NUPAGE 4-12% BT gels (Life Technologies, Darmstadt, Germany). For protein transfer, PROTRAN Western Blot nitrocellulose membranes (GE10600002; GE Healthcare, Breisgau, Germany) were used. Full original scans of the WBs are shown in Supplementary file. Antibody sources and dilutions are listed in the Supplementary Table S9.

Biotin was detected by incubation of the membrane with streptavidin-Alexa Fluor (1:10000, S21378, Thermo Fisher Scientific Inc. Waltham, USA) and then imaged using a LI-COR Odyssey CLx system.

### Immunoprecipitation

Wnt11 (Najdi *et al*, 2012), FZD6, Wnt11-V5, Wnt5a-V5 (MacDonald *et al*, 2014) were overexpressed in HEK293T cells for 48 hrs. Cells were collected by scratching and lysed as described above. Input samples were taken before addition of Protein G Dynabeads (Thermo Fisher, Darmstadt, Germany) with Wnt11/Ctrl antibody or V5 beads (A7345, Sigma, Sigma, St. Louis, MO, USA). In the indicated experiments, Wnt11 or Ctrl antibodies were cross-linked to the beads by BS3 (Thermo Fisher, Darmstadt, Germany) according to the manufacturer’s protocol, (30 min rotation at room temperature with 5 μM BS3 solution). Cell lysates were incubated with beads and the respective antibody for 18-22 hrs at 4 °C with rotation, beads were washed 4 times with PBS+1% Triton X-100 and once with PBS+0.1% Tween and eluted with 1x loading dye (in the case of FZD6 without DTT) at 70 °C for 10 min upon shaking. Antibody sources and dilutions are listed in Supplementary Table S9.

### Identification of acylation status of intracellular and secreted Wnt proteins

HEK293T wild type or Evi^KO^ were transfected with pcDNA Wnt3 or Wnt11 and labeled for 24 hrs with 50 µM clickable 9Z-hexadecenoic acid alkyne (palmitoleic acid alkyne, Alk-C16:1) (synthesized in multiple steps starting from 1,9-nonanediol according to procedure followed in (Tuladhar *et al*, 2019)) in DMEM with 10% dialyzed fetal bovine serum (Life Technologies, Grand Island, USA) was added.

Wnt11 from the supernatant was immunoprecipitated with Wnt11 antibody (GTX105971) and Pierce^TM^ Protein A/G magnetic beads (Thermo Fisher Scientific, USA). Then the click mix (1mM CuSO_4_ (Sigma-Aldrich, St. Louis, USA); 1mM TCEP (Carl Roth, Karlsruhe, Germany); 0.1mM Biotin-azide (Thermo Fisher Scientific Inc. Waltham, USA); 0.1mM TBTA) was added to the beads resuspended in PBS containing 0.05% Tween-20 in the ratio 1:10 for 1 hr. After the click reaction, the beads were washed three times with 0.05% Tween in PBS, each wash including a 10 min incubation of the beads in 0.05% Tween in PBS prior to centrifugation. The elution from the beads was performed by adding 100 µl of 2x Laemmli buffer and boiling for 5 min at 95 °C.

To investigate the palmitoleoylation status of intracellular Wnt proteins, cells were lysed and incubated with the click mix for 1 hr. Then ice-cold methanol, chloroform and water (4:1:3) were added to stop the reaction. After vortexing and centrifugation for 2 min at 16,000 g, 4°C upper phase was removed and ice-cold methanol (in ratio 1:2) was added. As the next step, vortexing and centrifugation were repeated to remove the methanol. The pellet was air dried for 10 min and solubilized in PBS with 4% SDS. After solubilization, 1% Triton X-100 in PBS was added to dilute the SDS concentration to 0.2%. Then a biotin pulldown using Pierce^TM^ Streptavidin Magnetic beads (Thermo Fisher Scientific Inc. Waltham, USA) was performed for 1hr at room temperature. After the pulldown, the beads were washed three times with 0.1% Tween in PBS, each wash including a 5 min incubation of the beads in 0.1% Tween in PBS prior to centrifugation. The elution from the beads was performed by adding 100 µl of 2x Laemmli buffer and boiling for 5 min at 95 °C.

### Luciferase assays

To measure the TCF4/Wnt-reporter activity, cells were seeded into 384-well white flat-bottom polystyrene plates (Greiner, Mannheim, Germany) (Voloshanenko *et al*, 2017). After 20-24 hrs, cells were transfected with 0.1 μl TransIT-LT1 transfection reagent (731-0029; VWR, Darmstadt, Germany), 20 ng TCF4/Wnt firefly luciferase reporter (Demir *et al*, 2013), 10 ng actin-*Renilla* luciferase reporter (Nickles *et al*, 2012), 20 ng Wnt3 (Najdi *et al*, 2012) or ctrl (pcDNA3)/β-catenin and 5 ng FZD6 (see also Supplementary Table S4). The amount of DNA was normalized by addition of a control plasmid (pcDNA3.1) for all conditions in the experiment. To assess paracrine induction of Wnt signaling, cells were treated with indicated amounts of recombinant mouse Wnt3a (PeproTech, Hamburg, Germany). Recombinant Wnt3a was added 16 hrs prior to the luciferase assay readout. Luminescence was measured with the Mithras LB940 plate reader (Berthold Technologies, Bad Wildbad, Germany). TCF4/Wnt-luciferase signal was normalized to the actin-*Renilla* luciferase reporter.

### Intestinal crypt organoid culture

The small intestine of Evi^fl/fl^ Cre_ERT (GPR177-ROSACre_ERT(tb3330)) mice (Augustin *et al*, 2013) was cut and processed for organoid formation as previously described (Mahe *et al*, 2013). Briefly, the mouse intestine was rinsed with ice-cold DPBS and villi were carefully scraped off using a metal spatula. Tissue fragments were incubated in 5 mM EDTA in DPBS on ice for 60 min and crypts were dissociated by vigorous shaking in the dissociation buffer. Finally, the crypt suspension was passed through a 70 μm filter. The flow-through was collected in a growth factor reduced matrigel and spotted onto suspension plates. Organoids were cultured in the medium described in the Supplementary Table S7. The medium was changed every second day and organoids were splitted every 6 to 7 days. For splitting, organoids were collected and centrifuged for 7-10 min at 4°C 300g. Medium was removed and 5 ml of cold DMEM + 10% FBS was added. For generation of Evi^KO^ organoids Evi^fl/fl^ Cre_ERT organoids were treated with 1 μM 4-HT (4-hydroxytamoxifen) for 4 days to deplete exon3 of *Evi/Wls*. For culturing Evi^KO^ organoids, 5 μM GSK3β inhibitor/CHIR99021 (361571, Merck-Millipore, Basel, Switzerland) was added to the medium because otherwise all Evi^KO^ organoids died within 4-5 days.

Matrigel was destroyed by repetitive pipetting using a 10 μl tip on top of a 10 ml serological pipet. Centrifugation was repeated and organoids were splitted in a ratio of 1:4. For further experiments (WB, FACS, IF) organoids were collected, centrifuged and incubated with 2-5 ml of cell recovery solution (354253; Corning, Wiesbaden) for 3-4 min for IF and for 6-9 min for WB and FACS. For detection of Wnt11 in the matrigel recovery solution was collected and diluted 1:3 with DMEM+10%FBS, 1% Triton-X100 and Blue Sepharose:PBS (1:1; 15 μl per 1 ml). Then the samples were treated as described for the Wnt secretion assay.

Animal welfare and experimental procedures were performed in accordance to German animal protection law (Permit number DKFZ223).

### Flow cytometry analysis (FACS)

HEK293T cells or single cells of organoids were washed with cold PBS and FACS buffer (PBS + 1% FBS). Organoid cells were incubated with the FcR Blocking Reagent (130-092-575; Miltenyi biotec, Bergisch Gladbach) for 30 min on ice. Then, samples were incubated with anti-Wnt11/V5/Ctrl antibody for 40-50 min on ice. Subsequently, cells were washed twice with FACS buffer and incubated with Alexa Fluor 647 goat anti-rabbit IgG (H+L) for 30 min on ice. After washing cells two more times with FACS buffer, Propidium Iodide (PI) solution was added for staining of dead cells. Measurements were done with the BD FACS Canto II. Single viable (PI negative) cells were gated for detection of Alexa Fluor 647. Representative gatings for the organoid experiments are shown in Supplementary file.

### Immunofluorescence of intestinal organoids

After 5 min of incubation in cell recovery solution, organoids were washed with cold PBS and fixed with 4% formaldehyde solution overnight at 4 °C. Then organoids were washed five times with cold PBS, incubated with 5% normal goat serum for 1 hour and stained with the primary antibody overnight. Then organoids were washed five times with cold PBS, stained with the secondary antibody for 1 hour at room temperature and washed five times with the cold PBS again. Immunofluorescence images were taken with a Leica TCS SP5 confocal microscope. Antibody sources and dilutions are listed in Supplementary table S8.

### Immunohistochemistry

Immunohistochemical analysis was performed on paraffin-embedded slides of Evi^fl/fl^ and Evi^KO^ mouse intestine as described in (Augustin *et al*, 2012). Antigen retrieval of paraffin sections was performed by microwaving the sections for 10 min in 10 mM citrate buffer, pH 6.0. The sections were incubated with the primary antibody overnight at 4°C according to the Supplementary Table S8. The secondary-biotinylated antibody (1:200, Dako, CA, USA) was incubated for 1 hr at room temperature followed by 30 min incubation with the Vectastain ABC-Kit (VEC-PK-6100, Biozol Diagnostica). Hematoxylin was used to stain the nuclei.

### Gelatin degradation assay

RPMI-7951 melanoma cells (150000) were transfected with 1.5 μg plasmid using FuGENE HD Transfection Reagent (E2311, Promega, Madison, USA.). 72 hrs later, cells were dissociated from the plate and seeded on gelatin-coated coverslips (30000 cells per 12 mm coverslip per condition). Gelatin coated slides were prepared using the QCM™ Gelatin Invadopodia Assay (Green) following the manufacturer’s instructions (ECM670, Merck Milipore, Basel, Switzerland). Additionally, Wnt protein expression in the lysate and supernatant was confirmed. 24 hrs after seeding on gelatin slides, cells were fixed using 4 % paraformaldehyde for 10 min at room temperature. For immunofluorescence staining of coverslips, cells were permeabilized using 0.2% Triton X-100/ PBS (v/v, Sigma-Aldrich, T8787) for 10 min and then blocked with blocking solution (3% FCS (v/v), 1% goat serum (v/v), 0.1% Triton X-100 (v/v) in PBS) for 30 min. Staining of the actin cytoskeleton was performed using Phalloidin-TRITC (dilution 1/500, provided with QCM™ Gelatin Invadopodia Assay (Green)) for 1 hr, followed by 3 washes in PBS. Coverslips were mounted using ProLong Diamond Antifade Mountant with DAPI (Life Technologies GmbH, P36962). Coverslips were imaged using a wide field Zeiss Cell Observer with a 20x objective. 8 randomly chosen fields of view with a minimum of 100 cells per condition were analyzed. Cell numbers were counted based on DAPI and actin staining. Images were analyzed using the Fiji (ImageJ) software. After applying the threshold function, the size of the black areas in each field of view was measured and this area was set in relation to the cell number. The capacity of gelatin degradation was determined per cell for each condition by normalizing to the respective control treatment for each replicate (values close to 1 indicate small difference between treatment and control). At least 6 biological replicates were performed per condition. The resulting values were tested against 1 using the Wilcoxon signed-rank test and software R, version 3.6.0, and R Studio.

### *Xenopus laevis* experiments

*In vitro* fertilization and embryo manipulation were performed as previously described (Gawantka *et al*, 1995). Developmental stages were determined according to Nieuwkoop and Faber (NF) (Nieuwkoop *et al*, 1994). NF stage 3 embryos were injected with *in vitro* transcribed mRNAs. The injected amount of mRNA ranged from 1 ng to 2 ng per embryo, as indicated in the respective figures. 5 nl was injected in the dorsal marginal zone of each blastomere and *ppl* was used as control mRNA. Quantifications were obtained by blind scoring of three independent experiments. All *X. laevis* experiments were executed according to federal and institutional guidelines and regulations and approved by the state review board of Baden-Württemberg, Germany (permit number G-141-18).

### ATF2 reporter assay

*X. laevis* embryos were injected at NF stage 3 with 1 ng to 2 ng of indicated mRNAs per embryo together with 200 pg *mFz* mRNA, 750 pg pRL-TK-Renilla and 200 pg ATF2 reporter plasmid. Embryo lysates were collected at NF stage 12 with 3 embryos per lysate. Dual luciferase reporter assay system (E1960, Promega, Madison, USA) was used for measurement of firefly (ATF2) and Renilla luminescence. Values for firefly luciferase were normalized to *Renilla* for every lysate. Statistical analyses were performed using unpaired t-test for 6 lysates from 3 independent injections.

### Modelling

Modelling was done using Alphafold 2.3.2 in multimer mode (Jumper *et al*, 2021) on bwForCluster Helix (date: 18.2.2025). Structural figures were prepared using ChimeraX-1.9 (Meng *et al*, 2023).

## ACKNOWLEDGEMENTS

We would like to acknowledge Kim Boonekamp, Antonia Schubert for critical comments on the manuscript and members of the Boutros and Bruegger groups for helpful discussions. We thank Dominique Dorr, Aileen Geib, Katrin Löffler and Frederika Reuter for help with experiments, and Jürgen Kopp (crystallization platform, BZH) for AlphaFold2 modeling support and helpful discussions. Furthermore, we are grateful for the support of the Core Facility for Mass Spectrometry & Proteomics (CFMP) at the ZMBH, Heidelberg University. We thank D. Virshup, M. Waterman and F. Zhang for plasmids which we received through Addgene. EJFM and IS acknowledge support by the state of Baden-Württemberg through bwHPC and the German Research Foundation (DFG) through grant INST 35/1597-1 FUGG and INST 35/1503-1 FUGG. This work was supported by the Deutsche Forschungsgemeinschaft (DFG, German Research Foundation) – project number 331351713 – SFB1324 on “Mechanisms and Functions of Wnt Signaling” and project number 278001972 – TRR/SFB 186 on “Molecular Switches in the Spatio-Temporal Control of Cellular Signal Transmission”.

## AUTHOR CONTRIBUTIONS

M.B., O.V., B.B., F.P. designed the study; O.V., N.D., D.A., A.I., L.W., D.K. performed experiments; I.A. provided Evi^fl/fl^ mice and made IHC; E.J.F.M. and I.S. performed modelling; C.S. and C.N. made *Xenopus* experiments; M.B., O.V., B.B., D.K., F.P. wrote the manuscript and all authors commented on the manuscript.

