## Supplemental Text for "Secretion and signaling properties of Wnt11 and Wnt3 are determined by their N-terminus"

**Supplementary Tables S1-S2**
**Supplementary Figures S1-S8**
**Supplementary Tables S3-S12**

| **HEK293T** | | | | | |
|---|---|---|---|---|---|
| Evi$^{KO2.1}$ | sgEvi_ex3 | Exon3 | Deletion 1 nt | Insertion 1 nt | |
| Evi$^{KO2.9}$ | sgEvi_ex3 | Exon3 | Deletion 1 nt<br>Deletion 2 nt | Deletion 26 nt | Insertion 1 nt |
| Evi$^{KO1.1}$ | sgEvi_ex2 | Exon2 | Deletion 2 nt<br>Deletion 4 nt | Insertion 1 nt | Deletion 9 nt<br>Deletion 3 nt |
| Evi$^{KO1.2}$ | sgEvi_ex2 | Exon2 | Deletion 2 nt | Deletion 5 nt<br>Mutation | Deletion 4 nt<br>Deletion 4 nt |
| **HCT116** | | | | | |
| Evi$^{KO2.1}$ | sgEvi_ex3 | Exon3 | Insertion 1 nt | | |
| Evi$^{KO2.2}$ | sgEvi_ex3 | Exon3 | Deletion 1 nt | Deletion 3 nt | |
| Evi$^{KO2.4}$ | sgEvi_ex3 | Exon3 | Deletion 12 nt | | |

**Supplementary Table 1. Mutations of *EVI/WLS* gene in knockout cells presented in this study.** DNA regions of indicated genes targeted by sgRNA were amplified by NestedPCR and then sequenced with the help of MiSeq. nt - nucleotide/s.

| | Unpaired *t* test with Welch's correction | | | | |
|---|---|---|---|---|---|
| | *p* (Ctrl *versus* Wnt11) | | | | |
| rec. Wnt3a | 0 | 12,5 | 25 | 50 | 100 |
| wt | 0.2061 | 0.0025 | 0.0005 | 0.0008 | 0.0002 |
| $Evi^{KO2.1}$ | 0.2952 | 0.0339 | 0.0777 | 0.1089 | 0.0053 |
| $Evi^{KO2.9}$ | 0.5947 | 0.0444 | 0.1515 | 0.133 | 0.0157 |
| | *p* (Ctrl *versus* Wnt5a) | | | | |
| rec. Wnt3a | 0 | 12,5 | 25 | 50 | 100 |
| wt | 0.7542 | 0.1004 | 0.0033 | <0.0001 | <0.0001 |
| Evi_2.1 | 0.4374 | 0.6034 | 0.8935 | 0.2036 | 0.6654 |
| Evi_2.9 | 0.4937 | 0.3997 | 0.3537 | 0.87 | 0.4496 |

**Supplementary Table 2. The table with *p* values for the Figure 1D, E.** *p* values were calculated with the help of GraphPad Prism10 by unpaired *t* test with Welch's correction.

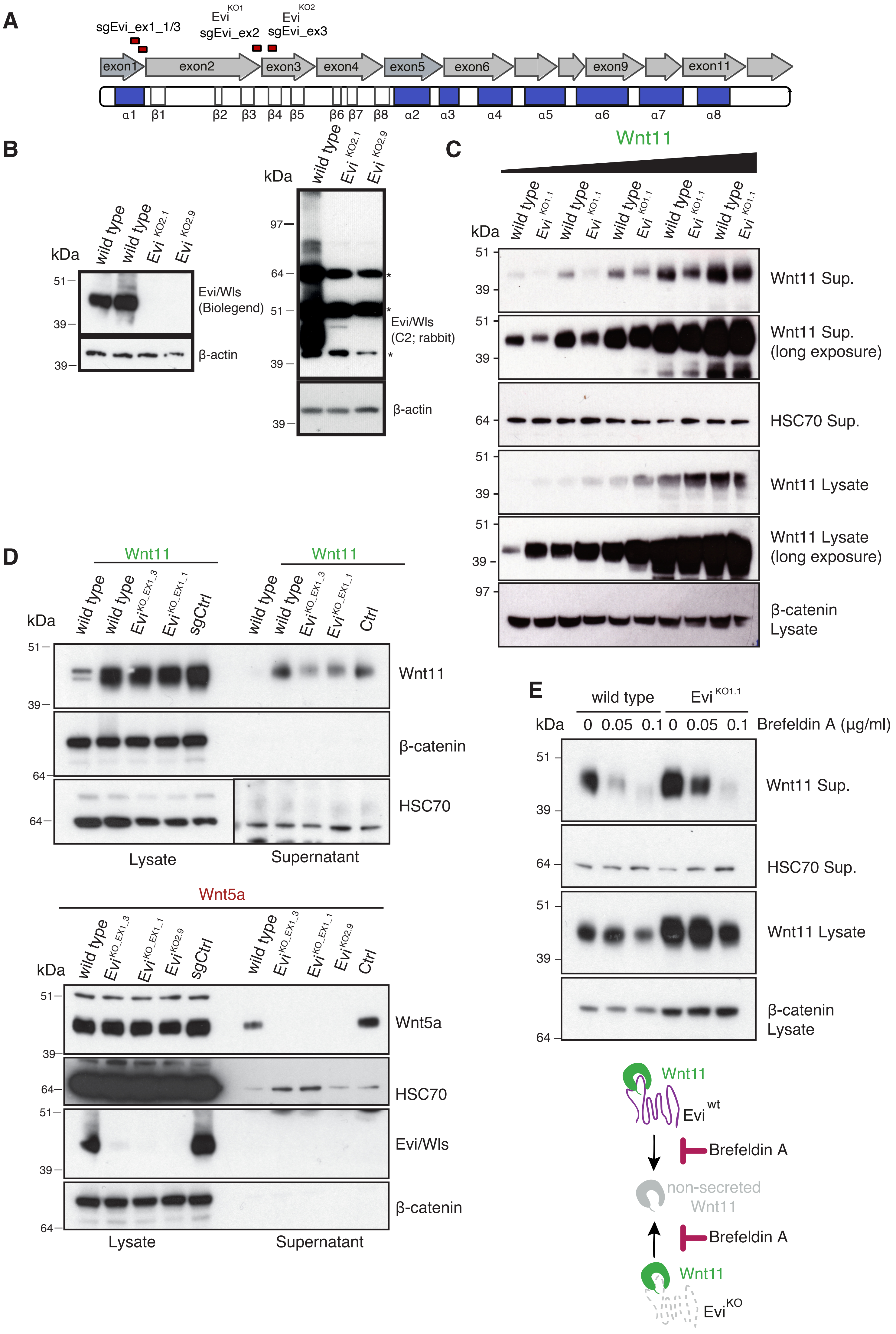

A
sgEvi_ex1_1/3
Evi KO1
sgEvi_ex2
Evi KO2
sgEvi_ex3
exon1
exon2
exon3
exon4
exon5
exon6
exon9
exon11
α1 β1 β2 β3 β4 β5 β6 β7 β8 α2 α3 α4 α5 α6 α7 α8
B
kDa
wild type
wild type
Evi KO2.1
Evi KO2.9
51
39
39
Evi/Wls (Biolegend)
β-actin
kDa
wild type
Evi KO2.1
Evi KO2.9
97
64
51
39
39
*
*
*
Evi/Wls (C2; rabbit)
β-actin
C
Wnt11
kDa
wild type
Evi KO1.1
wild type
Evi KO1.1
wild type
Evi KO1.1
wild type
Evi KO1.1
wild type
Evi KO1.1
51
39
Wnt11 Sup.
51
39
Wnt11 Sup. (long exposure)
64
HSC70 Sup.
51
39
Wnt11 Lysate
51
39
Wnt11 Lysate (long exposure)
97
β-catenin Lysate
D
Wnt11
Wnt11
kDa
wild type
wild type
Evi KO_EX1_3
Evi KO_EX1_1
sgCtrl
wild type
wild type
Evi KO_EX1_3
Evi KO_EX1_1
Ctrl
51
39
Wnt11
β-catenin
64
64
HSC70
Lysate
Supernatant
Wnt5a
kDa
wild type
Evi KO_EX1_3
Evi KO_EX1_1
Evi KO2.9
sgCtrl
wild type
Evi KO_EX1_3
Evi KO_EX1_1
Evi KO2.9
Ctrl
51
39
Wnt5a
64
HSC70
51
39
Evi/Wls
64
β-catenin
Lysate
Supernatant
E
wild type
Evi KO1.1
kDa
0 0.05 0.1 0 0.05 0.1
Brefeldin A (µg/ml)
51
39
Wnt11 Sup.
64
HSC70 Sup.
51
39
Wnt11 Lysate
64
β-catenin Lysate
Wnt11
Evi wt
Brefeldin A
non-secreted Wnt11
Brefeldin A
Wnt11
Evi KO

**Figure S1 Functional Wnt11 is secreted by  Evi knockout cells. A)** Schematic representation of the exon structure of Evi/Wls gene and corresponding domains of the protein. **B)** $Evi^{KO}$ HEK293T cells do not express Evi/Wls protein. Unspecific bands are marked with *. β-actin was used as the loading control. **C)** Less Wnt11 is secreted from $Evi^{KO}$ cells compared to wild type. $Evi^{KO1.1}$ cells, clone generated with sgEvi which targets exon2, were transfected with increasing amounts of the Wnt11 plasmid. **D)** Wnt11 secretion is not blocked in HEK293T $Evi^{KO}$ pools generated by sgRNA targeting exon 1, while Wnt5a secretion is blocked in these pools. **E)** Wnt11 secretion is blocked by Brefeldin A. $Evi^{KO1.1}$ or wild type HEK293T cells were transfected with Wnt11 and 24 hrs later they were treated with indicated amounts of Brefeldin A for 20 hrs. C-D) Wnts were precipitated from supernatant (Sup.) with Blue sepharose. β-catenin served as the loading control in the case of the lysates, HSC70 was used as the loading control for the supernatant.

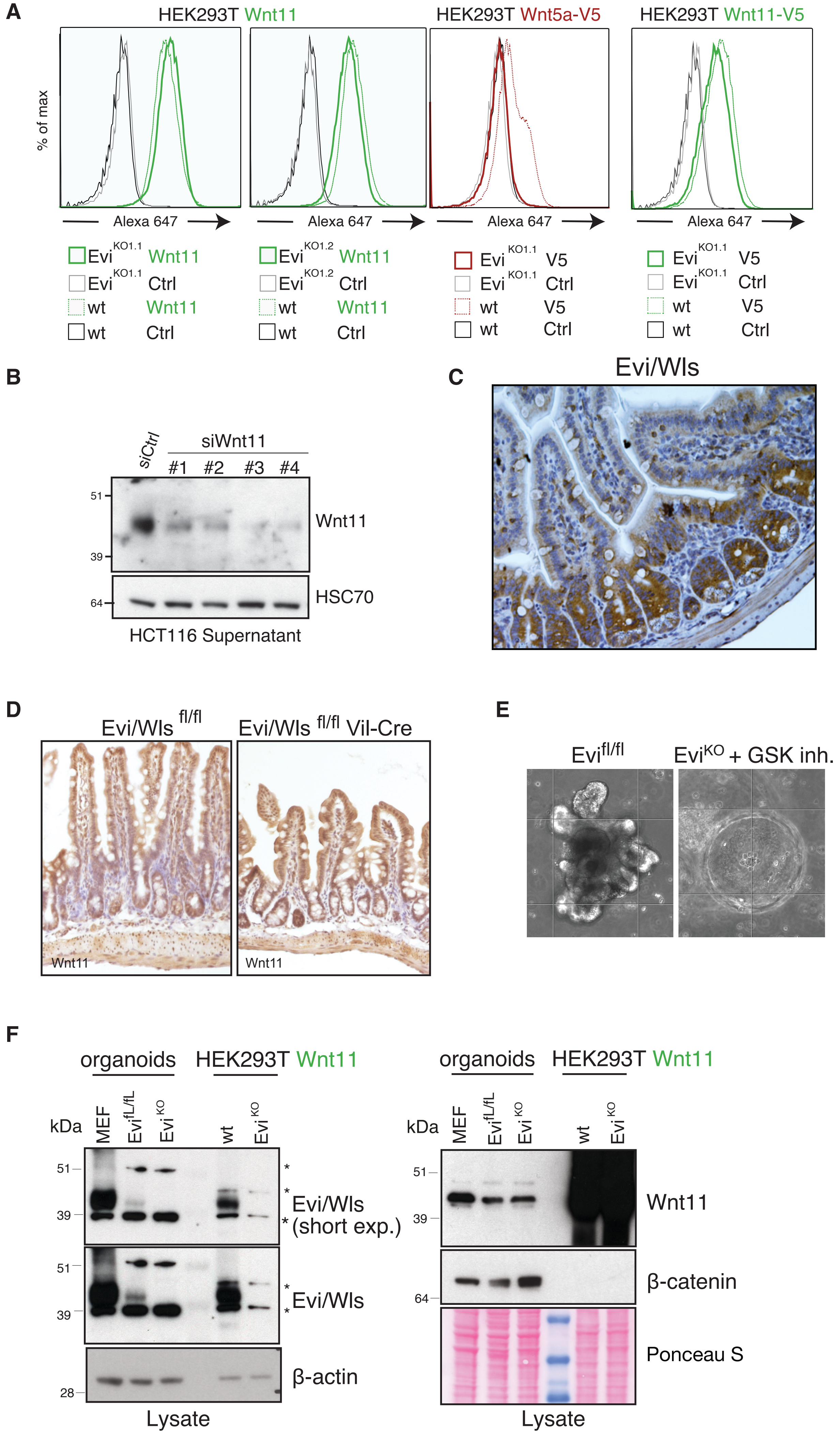

A
HEK293T Wnt11
HEK293T Wnt5a-V5
HEK293T Wnt11-V5
% of max
Alexa 647
Evi KO1.1 Wnt11
Evi KO1.1 Ctrl
wt Wnt11
wt Ctrl
Evi KO1.2 Wnt11
Evi KO1.2 Ctrl
Evi KO1.1 V5
Evi KO1.1 Ctrl
wt V5
wt Ctrl
B
siCtrl
siWnt11
#1 #2 #3 #4
51
39
64
Wnt11
HSC70
HCT116 Supernatant
C
Evi/Wls
D
Evi/Wls fl/fl
Evi/Wls fl/fl Vil-Cre
Wnt11
E
Evi fl/fl
Evi KO + GSK inh.
F
organoids
HEK293T Wnt11
kDa
MEF
Evi fL/fL
Evi KO
wt
51
39
28
64
Evi/Wls (short exp.)
Evi/Wls
β-actin
Lysate
Wnt11
β-catenin
Ponceau S

**Figure S2 Wnt11 is detected at the membrane of Evi$^{KO}$ intestinal organoids. A)** Wnt11 cell surface staining. Indicated HEK293T cells were transfected with the Wnt11, Wnt11-V5 or Wnt5a-V5 plasmids. 48 hrs later cells were stained with with primary antibodies against Wnt11 or V5. Then FACS analysis of Wnt11, Wnt11-V5 or Wnt5a-V5 in viable (PI-) wild type or Evi$^{KO}$ cells was performed. Representative results of 3 independent experiments are shown. **B)** Wnt11 antibody specificalli detects the endogenous protein. HCT116 cells were reverse transfected with indicated Wnt11 siRNAs for 96 hrs. After 48 hrs the medium was changed to remove old Wnt11. **C,D)** Evi/Wls and Wnt11 expression in mouse intestine. Immunohistochemical staining of Evi/Wls (B) or Wnt11 (C) was performed on the tissue slides of the small intestine. IHC slides were stained with anti-Evi(brown, C1; rabbit) or anti-Wnt11(brown,Abcam) antibody. Hematoxylin (blue) was used for the nucleus staining. **E)** Morphology of organoid cultures. Representative examples of organoids used in this study. **F)** Organoids show low expression levels of Evi/Wls. Mouse embryonic fibroblasts were used as control for detection of mouse Evi/Wls. β-actin served as the loading control.

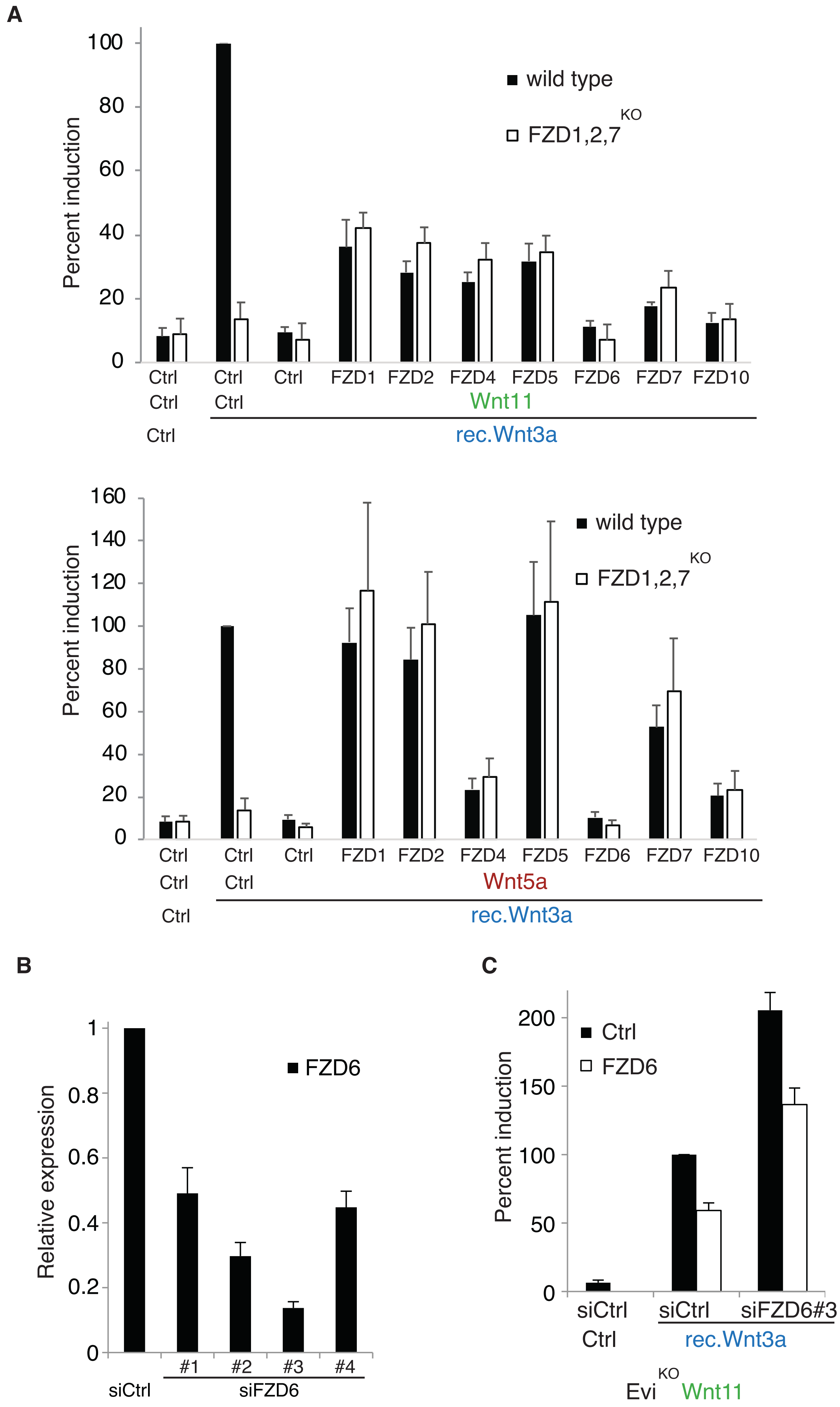
A
Percent induction
100
80
60
40
20
0
wild type
FZD1,2,7 KO
Ctrl Ctrl Ctrl FZD1 FZD2 FZD4 FZD5 FZD6 FZD7 FZD10
Ctrl Ctrl
Wnt11
Ctrl
rec.Wnt3a
Percent induction
160
140
120
100
80
60
40
20
0
wild type
FZD1,2,7 KO
Ctrl Ctrl Ctrl FZD1 FZD2 FZD4 FZD5 FZD6 FZD7 FZD10
Ctrl Ctrl
Wnt5a
Ctrl
rec.Wnt3a
B
Relative expression
1
0.8
0.6
0.4
0.2
0
FZD6
#1 #2 #3 #4
siCtrl
siFZD6
C
Percent induction
200
150
100
50
0
Ctrl
FZD6
siCtrl siCtrl siFZD6#3
Ctrl
rec.Wnt3a
Evi KO Wnt11

D

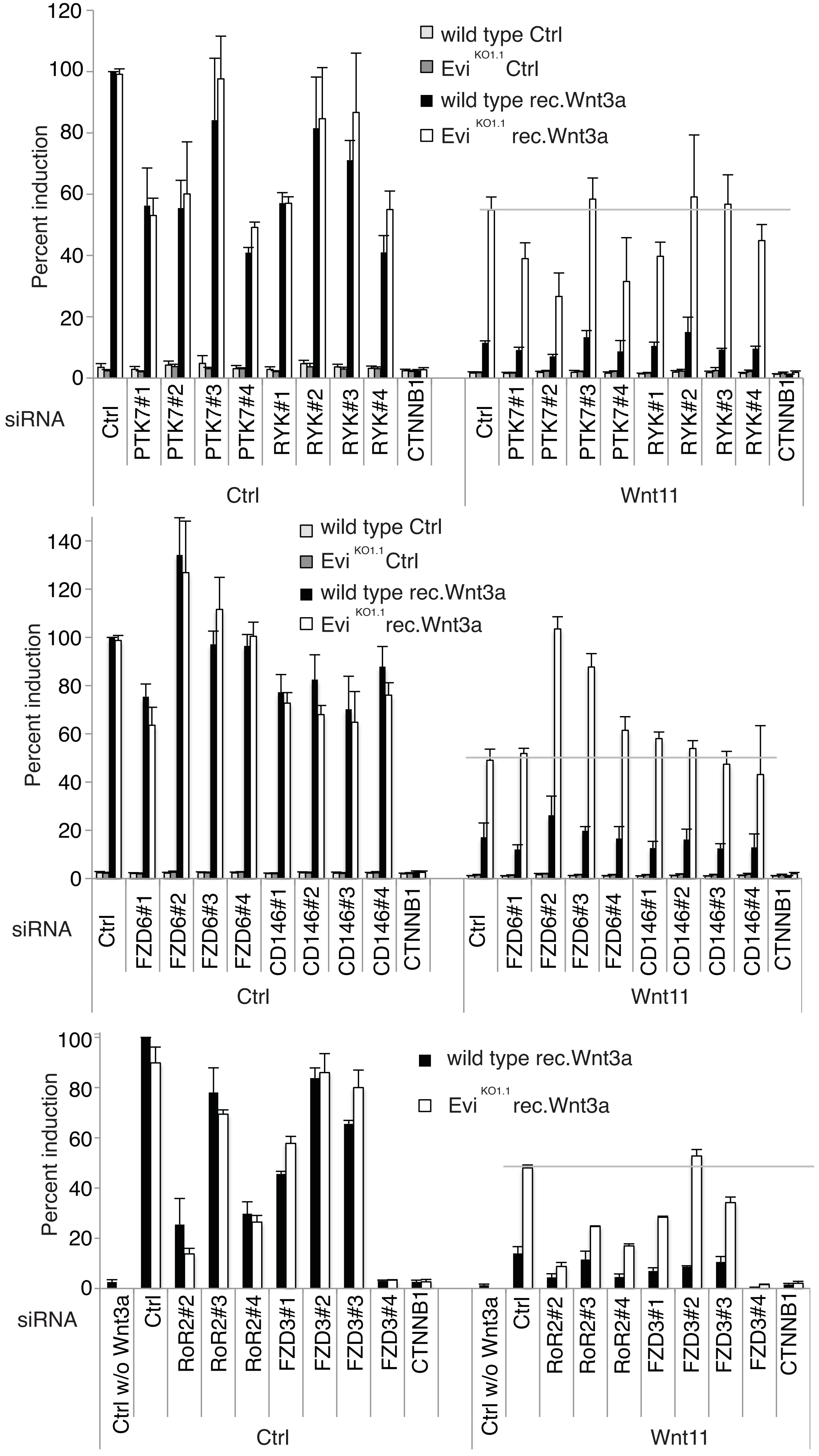

wild type Ctrl
Evi KO1.1 Ctrl
wild type rec.Wnt3a
Evi KO1.1 rec.Wnt3a
Percent induction
siRNA
Ctrl
PTK7#1
PTK7#2
PTK7#3
PTK7#4
RYK#1
RYK#2
RYK#3
RYK#4
CTNNB1
Ctrl
Wnt11
FZD6#1
FZD6#2
FZD6#3
FZD6#4
CD146#1
CD146#2
CD146#3
CD146#4
Ctrl w/o Wnt3a
RoR2#2
RoR2#3
RoR2#4
FZD3#1
FZD3#2
FZD3#3
FZD3#4

**Figure S3 Wnt11 secreted from wild type and Evi$^{KO}$ cells inhibits β-catenin-dependent Wnt signalling via the FZD6 receptor. A)** Overexpression of FZDs rescues Wnt3a-induced β-catenin-dependent signalling upon its inhibition with Wnt5a but not with Wnt11. **B)** mRNA levels of FZD6 after silencing with siRNAs. HEK293T Evi$^{KO}$ cells were reverse transfected for 72 hrs and then RT-qPCR was performed. **C)** Overexpression of FZD6 rescues siFZD6-induced phenotype of Wnt11 in Evi$^{KO}$ cells. Wnt11 stably transduced Evi$^{KO}$ cells were reverse transfected with indicated siRNAs. 24 hrs later the cells were transfected with FZD6 together with the TCF4/Wnt-firefly luciferase and actin-Renilla reporters for 48 hrs. 16 hrs before the read-out recombinant mouseWnt3a (100 ng/ml) was added. A-C) Results of 3(A,B)/4(C) independent experiments are shown as mean + SEM **D)** HEK293T wild type or Evi$^{KO}$ cells were reverse transfected with the indicated siRNAs for 72 hrs. Then the cells were transfected with Wnt11/Ctrl, TCF4/Wnt-firefly luciferase and actin-Renilla reporters for 48 hrs. 16 hrs before the read-out recombinant mouse Wnt3a (100 ng/ml) was added. Results of 3 independent experiments are shown as mean +SEM.

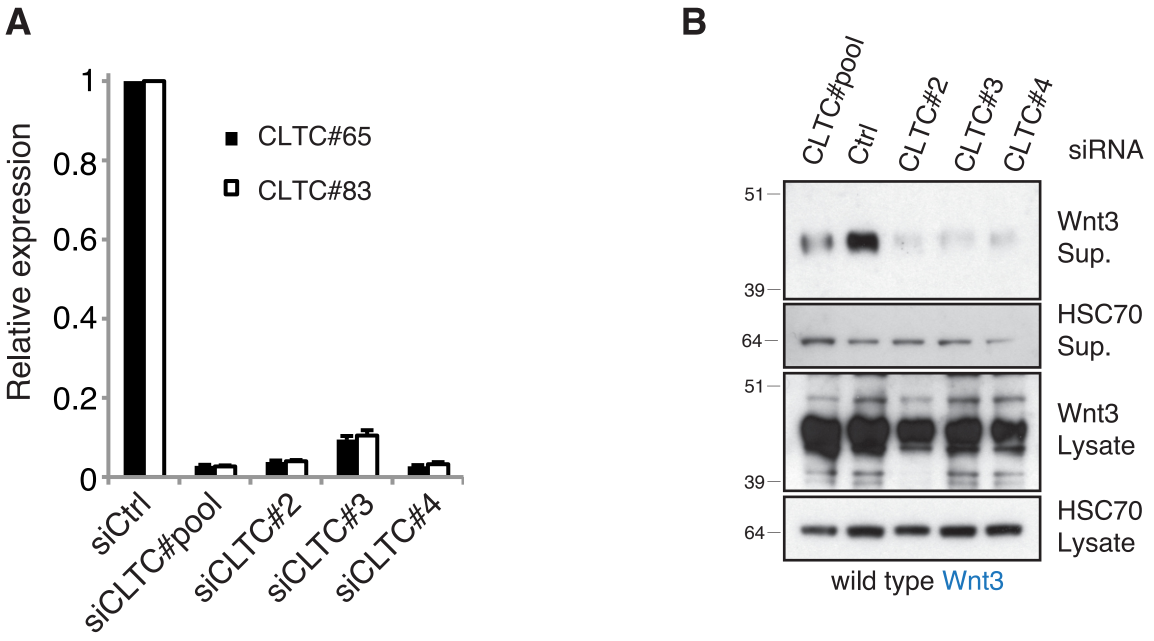

**Figure S4 Downregulation of clathrin inhibits secretion of Wnt3. A)** Silencing efficiency of siRNAs on mRNA level of clathrin heavy chain (CLTC). HEK293T cells were reverse transfected with siRNAs against CLTC for 72 hrs. Then RT-qPCR analysis was performed. Results are presented as mean +SEM of 3 independent experiments. **B)** HEK293T cells were reverse transfected with siRNAs against CLTC for 72 hrs and then transfected with Wnt3 plasmid for 48 hrs. Wnts were precipitated from supernatant (Sup.) with Blue sepharose. HSC70 served as the loading control.

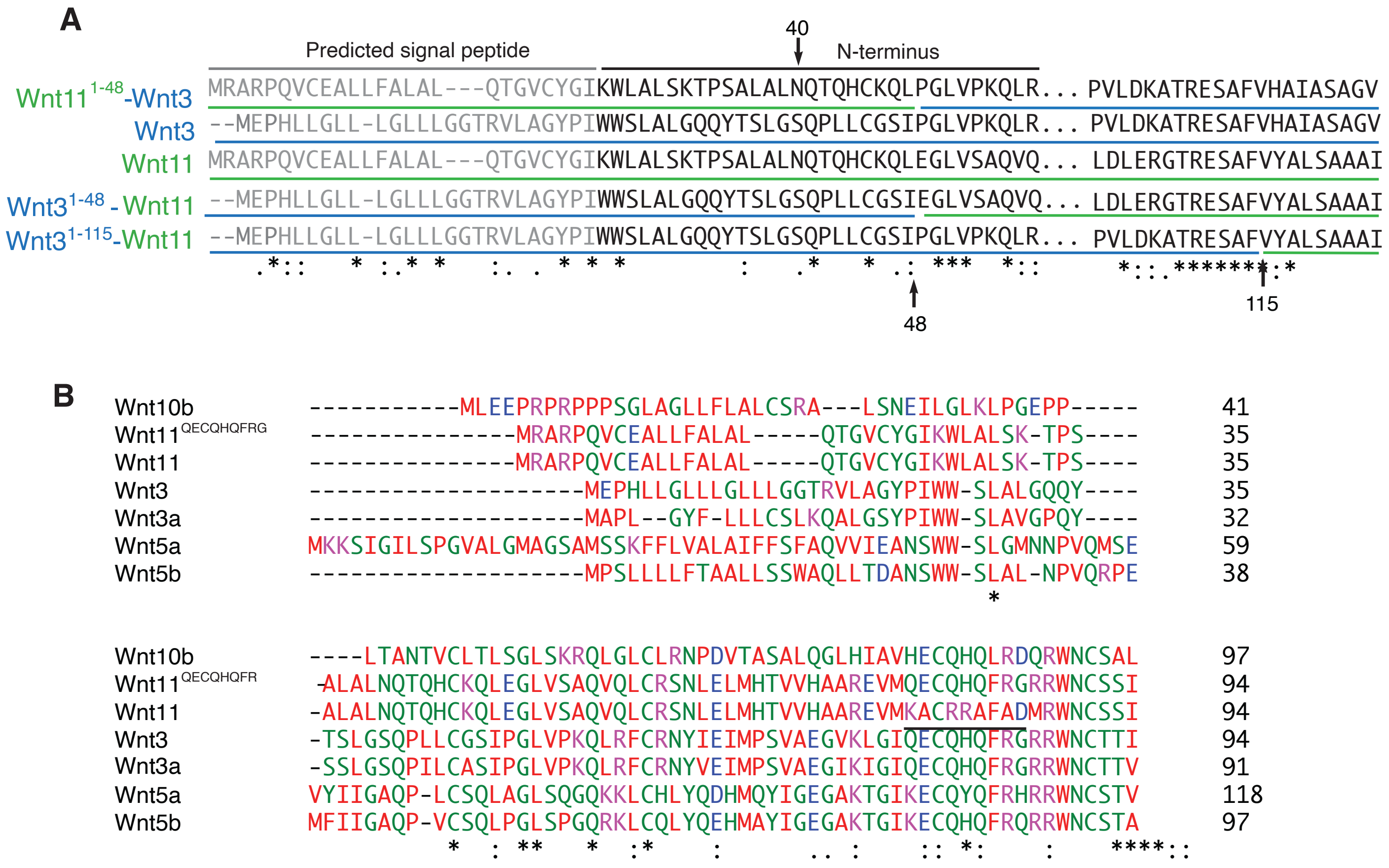

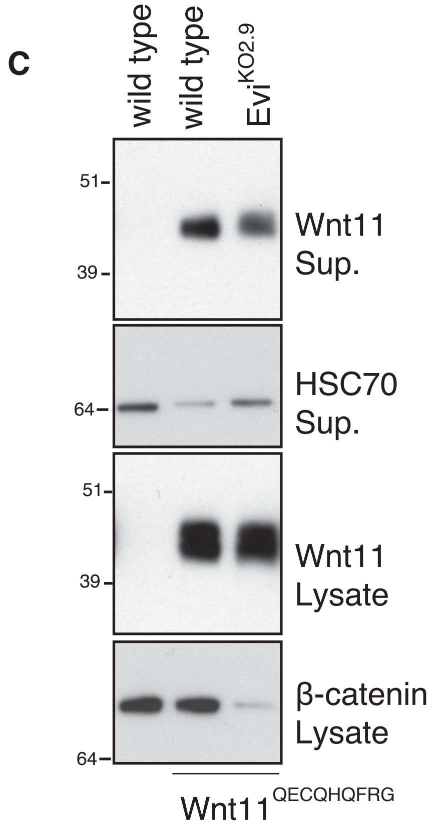

**Figure S5 Wnt11$^{QECQHQFRG}$ is still secreted in Evi$^{KO}$ cells. A,B)** Amino acid alignments of Wnt11, Wnt3, Wnt3$^{1\text{-}48}$-Wnt11, Wnt11$^{1\text{-}48}$-Wnt3 and Wnt3$^{1\text{-}115}$-Wnt11 proteins (A) and Wnt ligands (B). B) The underlined sequence in Wnt11 does not have homology to others Wnt ligands. **C)** Wnt11$^{QECQHQFRG}$ was expressed in wild type and Evi$^{KO}$ cells. HSC70 and β-catenin served as the loading controls. Representative results of 3 independent experiments are shown.

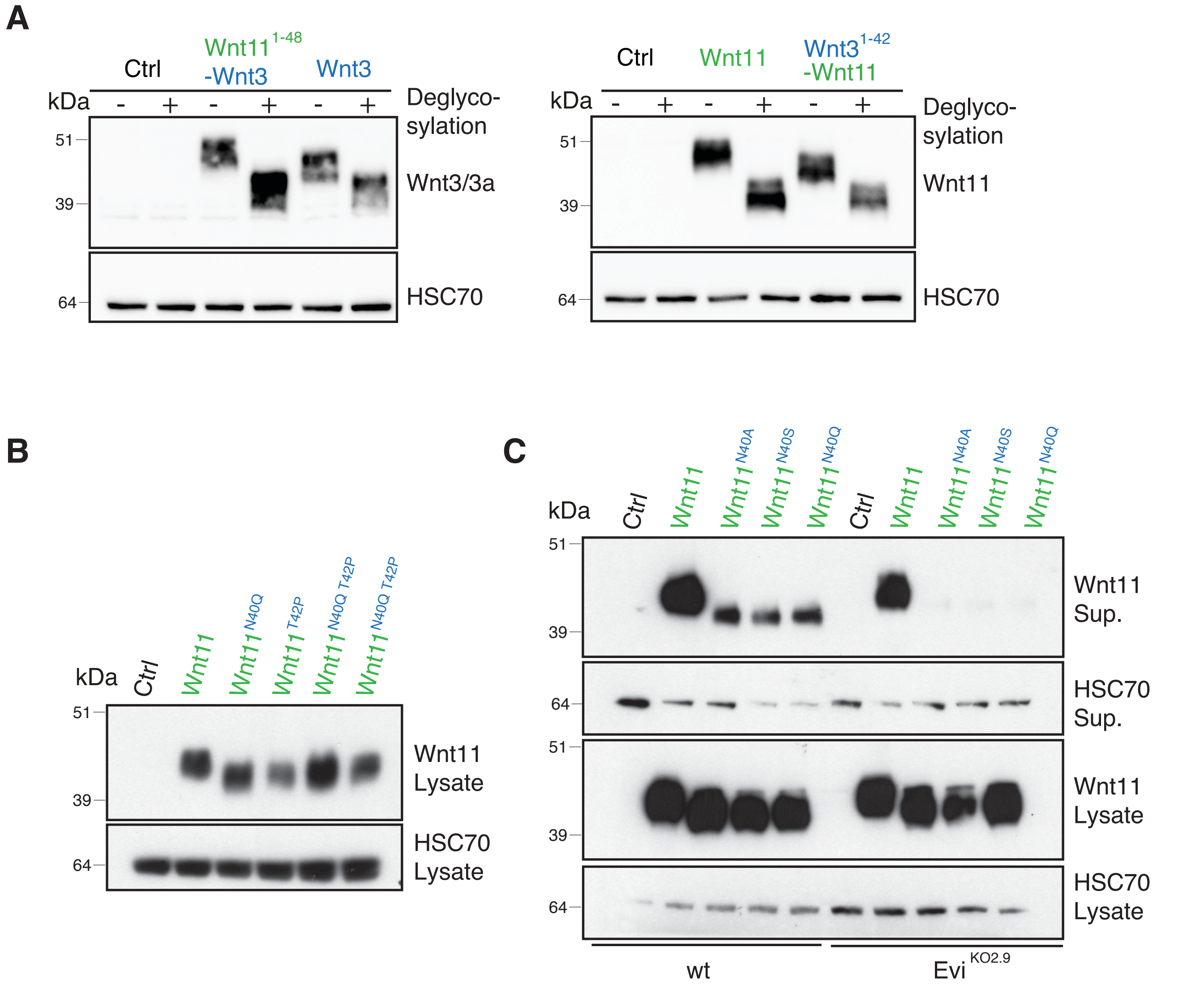

**Figure S6 Asparagine 40 of Wnt11 is glycosylated. A)** Wnt11 and Wnt3 show different glycosylation patterns in the N-terminus. Wnt3, Wnt11$^{1\text{-}48}$-Wnt3, Wnt11 and Wnt3$^{1\text{-}48}$-Wnt11 plasmids were transfected for 48 hrs and then the lysates were treated with deglycosylation mix of enzymes. **B)** Mutation of threonine 42 leads to deglycosylation of Wnt11 but does not enhance the phenotype induced by mutation of asparagine 40. **C)** Wnt11$^{N40A}$ and Wnt11$^{N40S}$ mutants show similar secretion behavior to Wnt11$^{N40Q}$ protein. HEK293T cells were transfected with the indicated Wnt11 mutants for 48 hrs. A-C) Representative results of 3 independent experiments are shown.

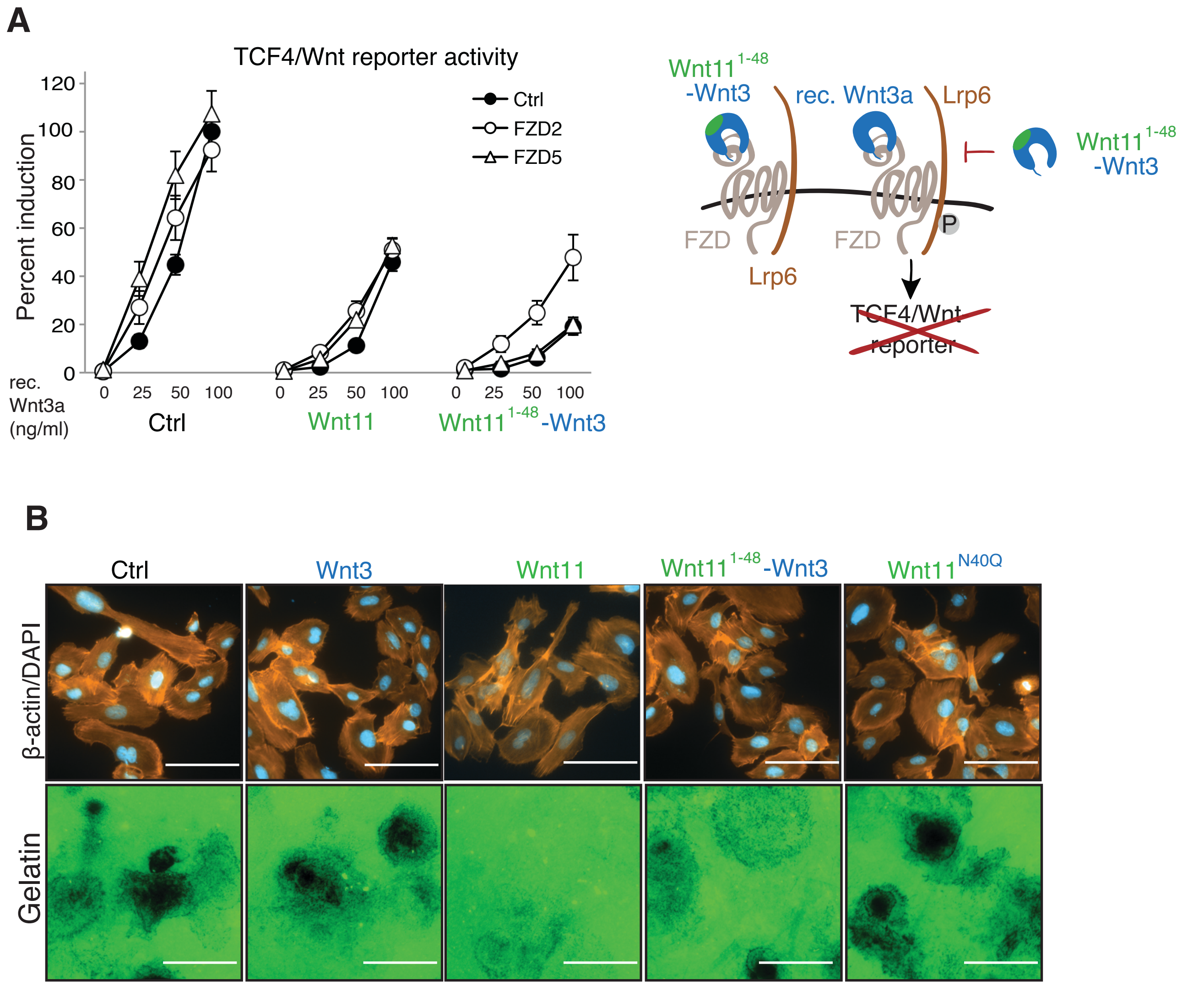

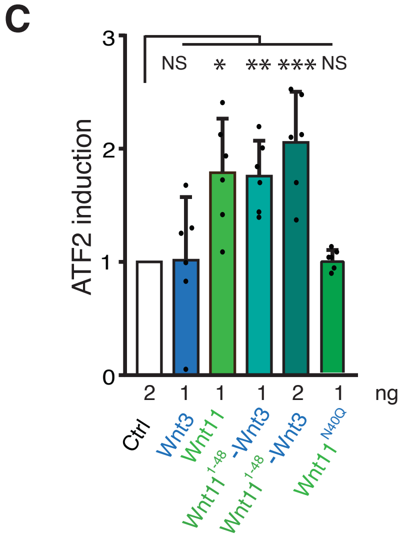

**Figure S7 The N-terminus of Wnt11 switches Wnt3 function from canonical to non-canonical. A)** FZD2 and FZD5 overexpression does not rescue Wnt11$^{1-48}$-Wnt3-induced inhibition of rec. Wnt3a-induced TCF4/Wnt-reporter activity. HEK293T were transfected with the indicated Ctrl, Wnt11$^{1-48}$-Wnt3, FZD2 or FZD5 constructs and with TCF4/Wnt-firefly luciferase and actin-Renilla reporters for 48 hrs. Recombinant mouse Wnt3a was added in the indicated amounts 16 hrs before the read-out. Results of 4 independent experiments are shown as mean +-SEM **B)** Wnt11 and Wnt11$^{1-48}$-Wnt3 reduce gelatin degradation capacity of RPMI-7951 melanoma cells. Cells were transfected with Ctrl, Wnt3, Wnt11, Wnt11$^{1-48}$-Wnt3 or Wnt11$^{N40Q}$ for 72 hrs. Then they were seeded on fluorescein-gelatin (green) coated coverslips for 24 hrs, later on fixed and stained for DNA (blue) and actin (orange). Scale bar is 100 µm. **C)** Wnt11$^{1-48}$-Wnt3 induces the ATF2 reporter in *Xenopus* embryos. Embryos were injected with the mFZD7 and indicated mRNAs, ATF2 firefly luciferase and TK-Renilla reporters at NF stage 3. The ATF2 reporter assay was performed at NF stage 12. Average +SD of 6 lysates (3 embryos per lysate) of 3 independent injections are shown. Every dot represents one lysate. Statistical significance, determined by the unpaired t-test, is represented by *p=0.0024, **p=0.0001, ***p=0.0002, NS, not significant.

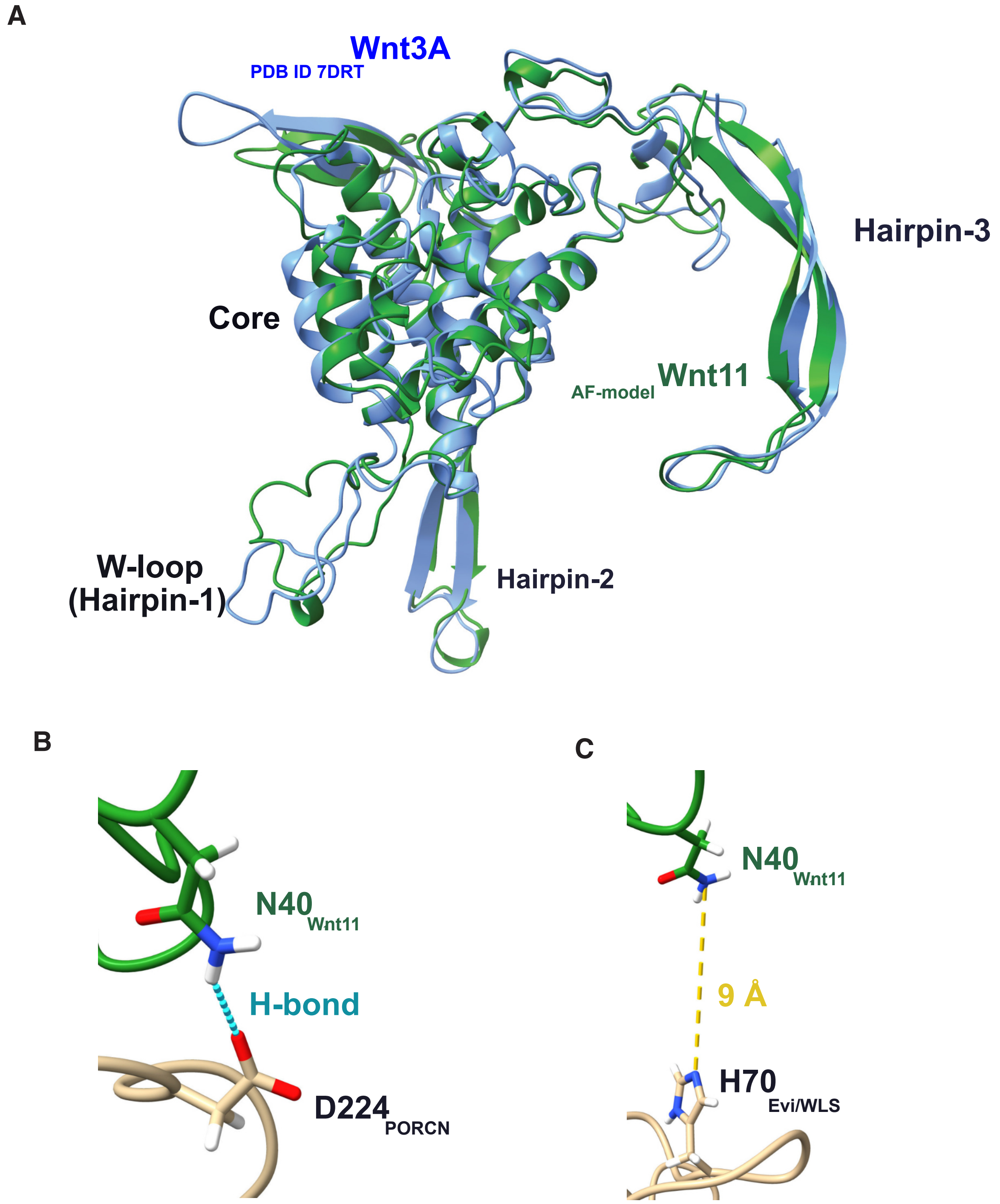

**Figure S8 Analysis of AlphaFold2 predictions. A)** Superposition of Wnt3A (blue; PDB ID 7DRT) with the AF2 prediction for Wnt11 (green). **B)** Detail of the PORCN-Wnt11 interaction model (AF2). $N40_{Wnt11}$ and $D224_{PORCN}$ are in H-bond distance. **C)** Detail of the Evi/WLS-Wnt11 interaction model (AF2). $N40_{Wnt11}$ and $H70_{WLS}$ are 9 Å apart.

**Supplementary Table S3**

**sgRNAs nucleotide sequences**

Notably, 5’-cacc-3’ and 5’-aaac-3’ are adapters required for cloning into the px459 vector backbone.

| sgRNA | KO | Sequence (5’-3’) |
|---|---|---|
| sgEvi_ex3 | KO2 | GTAAGCCAGGGAAACGTCCA |
| sgEvi_ex2 | KO1 | GAGAGTACACTTAACTGATT |
| sgEvi_ex1_1 | pools | gCAAATCATCGCCTTTCTGGT |
| sgEvi_ex1_3 | pools | gAATCAAGCCTCCCACCAGAA |

**Supplementary Table S4**

**Constructs**

| Short name | Name | Source |
|---|---|---|
| px459 | pSpCas9(BB)-2A-Puro (PX459) | Addgene #48139 - (Ran et al. 2013) |
| TCF4/Wnt luciferase reporter | 6xKD; pGL4.26 6xTcf-Firefly luciferase | K. Demir (Boutros lab, (Demir et al. 2013) |
| Renilla reporter/actin-*Renilla* | pAct-RL (Renilla luciferase) | D. Nickles (Boutros lab, (Nickles et al. 2012)) |
| Wnt11 | pcDNA3 Wnt11 | Addgene #35922- (Najdi et al. 2012) |
| Wnt11-V5 | pcDNA3 Wnt11-V5 | Addgene #43824- (Najdi et al. 2012) |
| Wnt11 | pCMV6-XL5 Wnt11 | Origene, SC303510 |
| Wnt3 | pcDNA3 Wnt3 | Addgene #35909 - (Najdi et al. 2012) |
| Wnt5a | pcDNA3 Wnt5a | Addgene # 35911 - (Najdi et al. 2012) |
| Wnt5a-V5 | pcDNA Wnt5a-V5 | Addgene #43813- (MacDonald et al. 2014) |
| Wnt11-V5 | pcDNA Wnt11-V5 | Addgene # 43824- (MacDonald et al. 2014) |

| | | |
|---|---|---|
| FZD1 | pCMV-XL4 FZD1 | Origene, SC117910 |
| FZD2 | pCMV-XL4 FZD2 | Origene, SC127603 |
| FZD4 | pCMV6-XL6 FZD4 | Origene, SC115479 |
| FZD5 | pcDNA3.2-FZD5-V5 | Boutros lab, re-cloned from pENTR223.1 FZD5 (DKFZ -Vector and Clone Repository of GPCF) |
| FZD6 | pCMV6-XL4 FZD6 | Origene, SC316995 |
| FZD7 | pCMV6-XL5 FZD7 | Origene, SC122259 |
| FZD10 | pCMV6-XL4 | Origene, SC115678 |
| β-catenin | pcDNA3 β-catenin | Boutros lab |
| Wnt11$^{N40Q}$ | Mutated pcDNA3 Wnt11 | Addgene #35922- (Najdi et al. 2012) |
| Wnt11$^{N40A}$ | Mutated pcDNA3 Wnt11 | Addgene #35922- (Najdi et al. 2012) |
| Wnt11$^{N40S}$ | Mutated pcDNA3 Wnt11 | Addgene #35922- (Najdi et al. 2012) |
| Wnt11$^{T42P}$ | Mutated pcDNA3 Wnt11 | Addgene #35922- (Najdi et al. 2012) |
| Wnt11 $^{N40Q, T42P}$ | Mutated pcDNA3 Wnt11 | Addgene #35922- (Najdi et al. 2012) |
| Wnt3$^{1-115}$-Wnt11 | pcDNA3 Wnt3$^{1-115}$-Wnt11 | Boutros lab |
| Wnt3$^{1-48}$-Wnt11 | pcDNA3 Wnt3$^{1-48}$-Wnt11 | Boutros lab |
| Wnt11$^{1-48}$-Wnt3 | pcDNA3 Wnt11$^{1-48}$-Wnt3 | Boutros lab |
| Wnt11 | pCS2+ Wnt11 | Boutros lab |
| Wnt3 | pCS2+ Wnt3 | Boutros lab |
| Wnt3$^{1-48}$-Wnt11 | pCS2+ Wnt3$^{1-48}$-Wnt11 | Boutros lab |
| Wnt11 $^{N40Q}$ | pCS2+ Wnt11 $^{N40Q}$ | Boutros lab |

**Supplementary Table S5**

**Primer adapters for Nested PCR 1**

| | |
|---|---|
| Forward adapter | 5'-TCCCTACACGACGctcttccgatct - gen specific primer sequence |

| | |
|---|---|
| Reverse adapter | 5'-AGTTCAGACGTGTGctcttccgatct - gen specific primer sequence |

**Supplementary Table S6**

**Primers used for Nested PCR 1 for validation of mutations generated by sgRNAs**

| sgRNA | Target gene | Forward primer (5'->3') | Reverse primer (5'->3') |
|---|---|---|---|
| sgEvi_ex3 | *EVI/WLS* | GTAAGCCAGGGAAACGTCCA | TGGACGTTTCCCTGGCTTAC |
| sgEvi_ex2 | *EVI/WLS* | ATGCCCGTAAGAACCATCAC | GGGAAGAAAGGGATGAGAGG |
| sgEvi_ex1 _1/3 | *EVI/WLS* | ATGGCTGGGGCAATTATAGA | TCATAGAAGCAAGGCAGTGA |
| | *EVI/WLS* | AATGGCTGGGGCAATTATAG | GATCATAGAAGCAAGGCAGTGA |

**Supplementary Table S7**

**Taqman qPCP primers and probes**

| Gene | Forward | Reverse | Probe |
|---|---|---|---|
| **FZD6** | gaagcaaaaagacatgcacaga | ttcgactttcactgattggatct | #23 |
| **CLTC** | ttccaaagaaggcagtggat | ccacatcatgcttttcactgat | #65 |
| **CLTC** | tttttgcccagggaaattact | ctggaaccgacggatagtgt | #83 |

**Supplementary Table S8**

**siRNAs**

| siRNA | siRNA ID Ambion/Dharmacon(D) | Sequence |
|---|---|---|
| siCtrl | | *Silencer*® Select Negative Control # 2 |
| siFZD6#1 | D-005505-01 | GGACAAGGATATAAGTTTC |
| siFZD6#2 | D-005505-03 | CACCAAACATTGAAACTTT |

| | | |
|---|---|---|
| siFZD6#3 | D-005505-04 | GCTTGAATGTGACAGATTA |
| siFZD#4 | D-005505-17 | CCTTAAATCATGTTCGACA |
| siWNT11#1 | D-009474-02 | CAGGATCCCAAGCCAATAA |
| siWNT11#2 | D-009474-03 | CGACAGCTGCGACCTTATG |
| siWNT11#3 | D-009474-04 | GTCGAGCGGTGCCACTGTA |
| siWNT11#4 | D-009474-05 | GGATCGGAACTCGTCTAT |

**Supplementary Table S9**

**Antibody**

| **Antibody** | **Reference** | **Dilution** | **Company** | **Source** |
|---|---|---|---|---|
| **Western blot** | | | | |
| Wnt11 | GTX105971 | 1:1000 | Genetex | Rabbit |
| Wnt11 | ab31962 | 1:1000 | Abcam | Rabbit |
| FZD6 (C12) | sc-393791 | 1:1000 | SantaCruz | Mouse |
| V5 | 600-401-378 | 1:1000 | Rockland | Rabbit |
| active-β-catenin (non-P-β-catenin) | 8E7/ 05-665 | 1:1000 | Millipore | Mouse |
| β-catenin | 610154 | 1:3000 | BD Bioscience | Mouse |
| HSC70 (B-6) | sc-7298 | 1:1000 | SantaCruz | Mouse |
| p-Lrp6 (Ser1490) | 2568 | 1:1000 | NEB | Rabbit |
| Lrp6 (C47E12) | 3395 | 1:1000 | NEB | Rabbit |
| Wnt3a/Wnt3 | ab28472 | 1:1000 | Abcam | Rabbit |
| Wnt3a | GTX128101 | 1:1000 | Genetex | Rabbit |
| β-actin (AC-40) | A3853 | 1:3000 | Sigma | Mouse |
| β-actin-HRP | sc-47778 | 1:10000 | SantaCruz | |
| Wnt5a/b (C27E8) | 2530 | 1:1000 | NEB | Rabbit |
| Evi | C2 | 1:500 | Boutros lab | Rabbit |
| Wls/Evi | 65590 | 1:1000 | BioLegend | Mouse |
| Anti-Mouse IgG (H+L) | 115-035-003 | 1:10000 | Jackson/Dianova | Goat |

| | | | | |
|---|---|---|---|---|
| Anti-Rabbit IgG (H+L) | 111-035-003 | 1:10000 | Jackson/Dianova | Goat |
| Streptavidin, Alexa Fluor™ 680 | S32358 | 1:10000 | ThermoFischer | |
| **Immunoprecipitation** | | | | |
| Wnt11 | GTX105971 | 1:700 | Genetex | Rabbit |
| Wnt3 | GTX128100 | 1:700 | Genetex | Rabbit |
| V5 beads | A7345 | 1:300-1:500 | Sigma | |
| VeriBlot for IP secondary antibody (HRP) | ab131366 | 1:3000 | Abcam | |
| **Immunofluorescence** | | | | |
| Wnt11 | GTX105971 | 1:1000 | Genetex | Rabbit |
| EpCAM- APC (CD326;G8.8) | 17-5791-80 | 1:500 | eBioscience | |
| Alexa Fluor® 488 goat anti-rabbit IgG (H+L) | A11078 | 1:500 | Invitrogen | |
| Alexa Fluor® 594 goat anti-mouse IgG (H+L) | A21203 | 1:500 | Invitrogen | |
| **FACS** | | | | |
| Wnt11 | GTX105971 | 1:100 | Genetex | Rabbit |
| Alexa Fluor® 647 goat anti-rabbit IgG (H+L) | A21245 | 1:200-1:400 | Invitrogen | |
| PI (propidium iodide) | BTIU40017 | 1 µg/ml | VWR | |
| **Immunochemistry** | | | | |
| Wnt11 | GTX105971 | 1:500 | Genetex | Rabbit |
| Evi | C1 | 1:500 | Boutros lab | Rabbit |

**Supplementary Table S10**

**Mediums and supplies**

| **Product** | **Dilution** | **Company** | **Reference** |
|---|---|---|---|
| Long term storage at +4 °C | | | |
| DMEM/F12 | Basic medium (500 ml) | Life Technologies | 12634028 |
| GlutaMAX (100x) | 1x (5 ml) | Life Technologies | 35050087 |
| Pen/Strep 100x | 1x (5 ml) | Life Technologies | 15140122 |
| HEPES | 10 µM (5 ml of 1M) | Sigma | H4034 |
| Upon addition of these components the medium was stored at +4 °C not longer then 10 days: | | | |
| mNoggin | 100ng/ml | Peprotech | 250-38-100 |
| B27 (50x) | 1x | Life Technologies | 17504044 |
| NAC | 1.25 mM | Sigma Aldrich | |
| R-spondin medium | 10% of medium | Produced from HEK293T R-spondin1 cells, Cultrex Trevigen | 3710-001-01 |
| mEGF | 50 ng/ml | Peprotech | 315-09-1000 |
| Y-27632 | 10 µM | Biozol Diagnostica | SEL-S1049 |
| Primocin | 100 µg/ml | InvivoGen | ant-pm-2 |
| Stored at -20 °C: | | | |
| Matrigel GFR | | VWR | 734-1101 |

### Supplementary Table S11

### Primers used mutagenesis

| Primers | Sequence (5'->3') |
|---|---|
| Wnt11$^{N40Q}$f | AAGACACCATCGGCCCTGGCACTGcaaCAGACGCAACACTGCAAGCAGCTGGA |
| Wnt11$^{N40Q}$r | TCCAGCTGCTTGCAGTGTTGCGTCTGttgCAGTGCCAGGGCCGATGGTGTCTT |
| Wnt11$^{N40Q,T42P}$f | AAGACACCATCGGCCCTGGCACTGcaaCAGccgCAACACTGCAAGCAGCTGGA |
| Wnt11$^{N40Q,T42P}$r | TCCAGCTGCTTGCAGTGTTGcggCTGttgCAGTGCCAGGGCCGATGGTGTCTT |
| Wnt11$^{N40S}$f | AAGACACCATCGGCCCTGGCACTGtcaCAGACGCAACACTGCAAGCAGCTGGAG |
| Wnt11$^{N40S}$r | CTCCAGCTGCTTGCAGTGTTGCGTCTGtgaCAGTGCCAGGGCCGATGGTGTCTT |
| Wnt11$^{N40A}$f | AAGACACCATCGGCCCTGGCACTGgcaCAGACGCAACACTGCAAGCAGCTGGA |
| Wnt11$^{N40A}$r | TCCAGCTGCTTGCAGTGTTGCGTCTGtgcCAGTGCCAGGGCCGATGGTGTCTT |
| Wnt11$^{T42P}$f | AAGACACCATCGGCCCTGGCACTGAACCAGcCGCAACACTGCAAGCAGCTGGA |
| Wnt11$^{T42P}$r | TCCAGCTGCTTGCAGTGTTGCGgCTGGTTCAGTGCCAGGGCCGATGGTGTCTT |

### Supplementary Table S12

### gBlocks

| gBlocks | Sequence (5'->3') |
|---|---|
| Wnt3$^{1-115}$-<br>Wnt11<br>(SnaBI-<br>BamHI) | TGGGACTTTCCTACTTGGCAGTACATC**TACGTA**TTAGTCATCGCTATTACCATG<br>GTGATGCGGTTTTGGCAGTACATCAATGGGCGTGGATAGCGGTTTGACTCACGG<br>GGATTTCCAAGTCTCCACCCCATTGACGTCAATGGGAGTTTGTTTTGGCACCAAA<br>ATCAACGGGACTTTCCAAAATGTCGTAACAACTCCGCCCCATTGACGCAAATGG<br>GCGGTAGGCGTGTACGGTGGGAGGTCTATATAAGCAGAGCTCTCTGGCTAACTA<br>GAGAACCCACTGCTTACTGGCTTATCGAAATTAATACGACTCACTATAGGGAGA<br>CCCAAGCTGGCTAGTTAAGCTATCAACAAGTTTGTACAAAAAAGCAGGCTCCGC<br>GGCCGCCCCCTTCACCATGGAGCCCCACCTGCTCGGGCTGCTCCTCGGCCTCCTG<br>CTCGGTGGCACCAGGGTCCTCGCTGGCTACCCAATTTGGTGGTCCCTGGCCCTGG<br>GCCAGCAGTACACATCTCTGGGCTCACAGCCCCTGCTCTGCGGCTCCATCCCAG<br>GCCTGGTCCCCAAGCAACTGCGCTTCTGCCGCAATTACATCGAGATCATGCCCA<br>GCGTGGCCGAGGGCGTGAAGCTGGGCATCCAGGAGTGCCAGCACCAGTTCCGG<br>GGCCGCCGCTGGAACTGCACCACCATAGATGACAGCCTGGCCATCTTTGGGCCC<br>GTCCTCGACAAAGCCACCCGCGAGtctGCCTTCGTGTATGCGCTGTCGGCCGCCGC<br>CATCAGCCACGCCATCGCCCGGGCCTGCACCTCCGGCGACCTGCCCGGCTGCTC<br>CTGCGGCCCCGTCCCAGGTGAGCCACCCGGGCCCGGGAACCGCTGGGGAGGAT<br>GTGCGGACAACCTCAGCTACGGGCTCCTCATGGGGGCCAAGTTTTCCGATGCTC<br>CTATGAAGGTGAAAAAAACA**GGATCC**CAAGCCAATAAACTGATGCGTCTACA<br>CAACAGTGAAGTGGGGAGACAGGCT |
| Wnt3$^{1-48}$-<br>Wnt11<br>(XhoI and | GGCTT**cTCGAg**ATTAATACGACTCACTATAGGGAGACCCAAGCTtGCTAGTTAA<br>GCTATCAACAAGTTTGTACAAAAAAGCAGGCTCCGCGGCCGCCCCCTTCACCAT<br>GGAGCCCCACCTGCTCGGGCTGCTCCTCGGCCTCCTGCTCGGTGGCACCAGGGT<br>CCTCGCTGGCTACCCAATTTGGTGGTCCCTGGCCCTGGGCCAGCAGTACACATCT<br>CTGGGCTCACAGCCCCTGCTCTGCGGCTCCATCGAGGGTCTGGTGTCTGCACAG |

| | |
|---|---|
| KflI) | GTGCAGCTGTGCCGCAGCAACCTGGAGCTCATGCACACGGTGGTGCACGCCGCC<br>CGCGAGGTCATGAAGGCCTGTCGCCGGGCCTTTGCCGACATGCGCTGGAACTGC<br>TCCTCCATTGAGCTCGCCCCCAACTATTTGCTTGACCTGGAGAGA**GGGACCC**G<br>GGAG |
| Wnt11$^{1-48}$-Wnt3 (NotI and SfiI) | AGCTGGCTAGTTAAGCTATCAACAAGTTTGTACAAAAAAGCAGGCTCC**GCGGC**<br>**CGC**CCCCTTCACCATGAGGGCtCGGCCGCAGGTCTGCGAGGCGCTGCTCTTCGC<br>CCTGGCGCTCCAGACCGGCGTGTGCTATGGCATCAAGTGGCTGGCGCTGTCCAA<br>GACACCATCGGCCCTGGCACTGAACCAGACGCAACACTGCAAGCAGCTGCCAG<br>GCCTGGTCCCCAAGCAACTGCGCTTCTGCCGCAATTACATCGAGATCATGCCCA<br>GCGTGGCCGAGGGCGTGAAGCTGGGCATCCAGGAGTGCCAGCACCAGTTCCGG<br>GGCCGCCGCTGGAACTGCACCACCATAGATGACAGCCTGGCCATCTTTGGGCCC<br>GTCCTCGACAAAGCCACCCGCGAGTCGGCCTTCGTTCACGCCATCGCCTC**GGCC**<br>**GGCGTGGCC**TTCGCCGTCACCCGCTCCTGCGCCGAGGGCACCTCCACCATTTG<br>CGGCTGTGACTCGCATCA |
| Wnt11$^{QECQHQFRG}$ (SnaBI-BamHI) | TGGGACTTTCCTACTTGGCAGTACATC**TACGTA**TTAGTCATCGCTATTACCATG<br>GTGATGCGGTTTTGGCAGTACATCAATGGGCGTGGATAGCGGTTTGACTCACGG<br>GGATTTCCAAGTCTCCACCCCATTGACGTCAATGGGAGTTTGTTTTGGCACCAAA<br>ATCAACGGGACTTTCCAAAATGTCGTAACAACTCCGCCCCATTGACGCAAATGG<br>GCGGTAGGCGTGTACGGTGGGAGGTCTATATAAGCAGAGCTCTCTGGCTAACTA<br>GAGAACCCACTGCTTACTGGCTTATCGAAATTAATACGACTCACTATAGGGAGA<br>CCCAAGCTGGCTAGTTAAGCTATCAACAAGTTTGTACAAAAAAGCAGGCTCCGC<br>GGCCGCCCCCTTCACCATGAGGGCGCGGCCGCAGGTCTGCGAGGCGCTGCTCTT<br>CGCCCTGGCGCTCCAGACCGGCGTGTGCTATGGCATCAAGTGGCTGGCGCTGTC<br>CAAGACACCATCGGCCCTGGCACTGAACCAGACGCAACACTGCAAGCAGCTGG<br>AGGGTCTGGTGTCTGCACAGGTGCAGCTGTGCCGCAGCAACCTGGAGCTCATGC<br>ACACGGTGGTGCACGCCGCCCGCGAGGTCATGAAGGaaTGTCaaCatcaaTTTcggggcc<br>gtCGCTGGAACTGCTCCTCCATTGAGCTCGCCCCCAACTATTTGCTTGACCTGGAG<br>AGAGGGACCCGGGAGTCGGCCTTCGTGTATGCGCTGTCGGCCGCCGCCATCAGC<br>CACGCCATCGCCCGGGCCTGCACCTCCGGCGACCTGCCCGGCTGCTCCTGCGGC<br>CCCGTCCCAGGTGAGCCACCCGGGCCCGGGAACCGCTGGGGAGGATGTGCGGA<br>CAACCTCAGCTACGGGCTCCTCATGGGGGCCAAGTTTTCCGATGCTCCTATGAA<br>GGTGAAAAAAACA**GGATCC**CAAGCCAATAAACTGATGCGTCTACACAACAGT<br>GAAGTGGGGAGACAGGCTC |
